# Structural mechanism of lipid modulation of pentameric ligand-gated ion channel activity

**DOI:** 10.1101/2025.10.07.680764

**Authors:** Brandon K. Tan, Hanrui Xu, Jesse W. Sandberg, Grace Brannigan, Wayland W. L. Cheng

## Abstract

Pentameric ligand-gated ion channels (pLGICs) are sensitive to the lipid environment. However, the structural mechanism of how specific lipids support the agonist response of any pLGIC is poorly understood. Using the model pLGIC, ELIC (*Erwinia* ligand-gated ion channel), we find that phosphatidylethanolamine (PE) or cardiolipin (CL) are sufficient to support activation of a non-desensitizing mutant called ELIC5. Cryo-EM structures of unliganded and agonist-bound ELIC5 in the absence of PE or CL show increased structural heterogeneity and destabilization of the resting and open-channel states. Importantly, the unliganded structure of ELIC5 in a phosphatidylcholine (PC)-only environment shows variability that resembles agonist-induced changes. The structures also reveal a CL binding site at an outer leaflet M3-M4 site. Together with functional measurements in asymmetric liposomes and coarse-grained molecular dynamics simulations, the data indicate that CL supports ELIC activity by binding to this M3-M4 site thereby stabilizing an agonist-responsive resting state of the channel.

## Introduction

Pentameric ligand-gated ion channels (pLGICs), including the nicotinic acetylcholine receptor (nAChR) and GABA(A) receptor (GABA_A_R), are the principal mediators of synaptic transmission and neuronal excitability. These ion channels respond to or are activated by agonists like acetylcholine and GABA. They are also sensitive to their lipid environment (1), and there is increasing evidence that the function of these channels is modulated by their distribution in lipid microdomains (2–4). The nAChR requires certain phospholipids (phosphatidylethanolamine and phosphatidylserine) and cholesterol to support agonist response or ion channel activity (5–7), and the GABA_A_R and serotonin receptor (5-HT3_A_R) may also be modulated by cholesterol (4, 8, 9). However, the structural mechanism of lipid modulation of pLGIC function is poorly understood. Moreover, cryo-EM structures of many pLGICs in lipid nanodiscs or liposomes show phospholipid-like densities associated with the transmembrane domain (TMD) (10, 11), but it is unclear the extent to which phospholipid interactions at these sites influence pLGIC function.

The prokaryotic pLGIC, ELIC (*Erwinia* ligand-gated ion channel), remains an indispensable model for dissecting lipid modulation of this family of ion channels because of the ease of reconstituting ELIC into liposomes for functional studies (12–14). Earlier work established that the anionic phospholipid, phosphatidylglycerol (PG), decreases ELIC desensitization (13, 14), and structural and computational data support a mechanism whereby PG stabilizes the open-channel state relative to the desensitized state by binding to an outer leaflet site formed by the M4, M3 and M1 helices (13). It was also shown that maximal agonist responses in ELIC require both PG and phosphatidylethanolamine (PE) (13). This is reminiscent of the lipid requirements in the nAChR: maximal agonist response, as measured by ion fluxes, requires an anionic phospholipid (PS) and zwitterionic phospholipid, PE (5). Therefore, understanding the mechanism by which PE or other phospholipids support agonist responses of ELIC could provide insight into lipid modulation of other pLGICs.

In this study, we establish the phospholipids required to support ELIC agonist response and describe a structural mechanism. Using ELIC5—a mutant that does not desensitize—we find that PE or cardiolipin (CL) are sufficient to support ELIC activation. Cryo-EM structures of ELIC5 in lipid nanodiscs indicate that PE and CL stabilize the resting and open-channel structures of ELIC in the absence or presence of agonist, respectively. The structures without these lipids display increased conformational heterogeneity, and notably the unliganded structures show changes resembling partial activation. The structures also reveal bound CL in an outer leaflet site adjacent to the M3 and M4 helices. Using coarse-grained molecular dynamics (CGMD) simulations and functional measurements in asymmetric liposomes, the data suggest that CL supports ELIC5 agonist response by binding to an outer leaflet site and stabilizing the agonist-responsive resting state of the channel.

## Results

### CL recapitulates the effects of PG and PE on ELIC agonist response

We used a stopped-flow thallium flux assay to characterize ELIC activity in liposomes (Figure 1A). With the application of agonist, ELIC rapidly activates reaching a peak activity, and then slowly desensitizes. The peak activity or peak thallium flux rate of ELIC in response to agonist (henceforth called the peak agonist response) was significantly higher in liposomes containing a ternary mixture (2:1:1 molar ratio of POPC/POPG/POPE) compared to POPC alone (13). This difference in peak agonist response in the two lipid conditions could arise from a difference in ELIC channel activity, or a difference in the number of outwardly-facing channels per liposome. Only channels with the extracellular domain, which contains the agonist binding site, facing the extra-liposomal space are activated in the assay. To address this possibility, we quantified outwardly-facing channels in each liposome preparation (15). On a cysteine-less ELIC background (C300S/C313S), a cysteine was introduced to the N-terminus (A6C) and the detergent purified protein modified with the maleimide fluorophore, DY647P1. After reconstitution in liposomes, the quantity of outwardly-facing channels was determined by measuring the fluorescence intensity before and after treatment with TCEP, which quenches DY647P1 in the extra-liposomal space (Supplementary Figure 1A). DY647P1-labeled ELIC (A6C/C300S/C313S) showed ∼10x higher peak agonist response in 2:1:1 POPC:POPE:POPG liposomes compared to POPC liposomes (Fig. 1B) (13). However, the quantity of outwardly-facing channels in 2:1:1 POPC:POPE:POPG liposomes was only ∼1.6x higher than in POPC liposomes, meaning that the higher peak agonist response in the ternary lipid mixture is mostly attributed to higher channel activity (Supplementary Fig. 1B and 1C).

**Figure 1:**
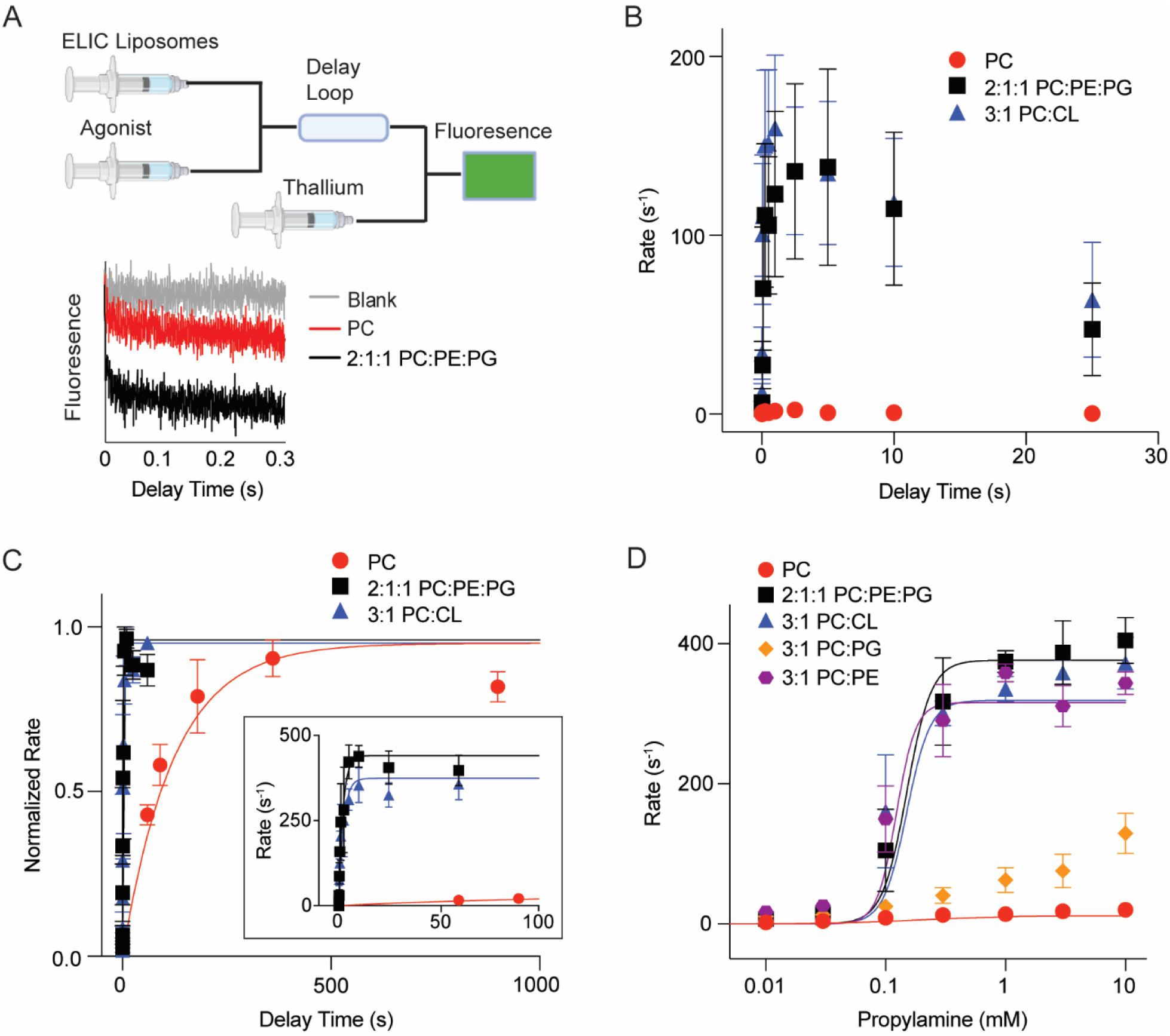
Effects of POPE/POPG and DPOCL on ELIC peak agonist response and desensitization: (A) *Top:* Schematic of the sequential mixing experiment for the fluorescence liposomal stopped-flow assay (created with BioRender.com). ELIC proteoliposomes are mixed with agonist; after a variable delay time, the sample is mixed with thallium, and the rate of fluorescence quenching is determined. *Bottom*: Representative fluorescence traces of ELIC5 in 2:1:1 liposomes without agonist (blank) or ELIC5 in POPC or 2:1:1 liposomes after mixing with 10 mM propylamine. (B) Tl^+^ flux rates of ELIC (A6C/C300S/C313S-labeled with DY647P1) in 2:1:1 POPC:POPE:POPG (black), 3:1 POPC:DPOCL (blue) and POPC (red) liposomes as a function of time after mixing with 10 mM propylamine (n = 3-8). (C) Normalized Tl^+^ flux rates of ELIC5 in POPC, 2:1:1 POPC:POPE:POPG, and 3:1 POPC:DPOCL liposomes (n = 3-6). The inset shows absolute Tl^+^ flux rates. (D) Tl^+^ flux rates of ELIC5 in liposomes with the indicated lipid compositions. Measurements were taken after a 20 min pre-incubation with varying propylamine concentration (n = 3-6). Data are fit to a Hill equation. All data are shown as mean ± SEM for (n) independent experiments.

A previously-reported cryo-EM structure of ELIC extracted from *e. coli* membranes using styrene maleic acid showed bound cardiolipin (CL) (16). Therefore, we examined the effect of CL on ELIC activation and desensitization using the stopped-flow thallium flux assay. Fluorescently-labeled ELIC (A6C/C300S/C313S) reconstituted in 3:1 POPC:DPOCL (di-16:0-18:1 cardiolipin) liposomes showed higher peak agonist response and slower desensitization compared to POPC liposomes (Figure 1B, Supplementary Figure 1D), and the difference in peak agonist response is not accounted for by the quantity of outwardly-facing channels (Supplementary Fig. 1B and 1C). Therefore, DPOCL recapitulates the dual effects of POPG and POPE on ELIC function: like POPG, it slows desensitization, and like POPG and POPE, it produces a higher peak agonist response.

### PE or CL are sufficient to support ELIC activation

The effect of POPG/POPE or DPOCL on peak agonist response could arise from an effect on activation or desensitization. For example, rapid and profound desensitization of ELIC in POPC liposomes could account for the low peak agonist response. To examine the effect of lipids on ELIC activation without the confounding influence of desensitization, we analyzed ELIC5, a combination of five mutations (P254G/C300S/V261Y/G319F/I320F) in ELIC that eliminate channel desensitization (13). Activation rates of ELIC5 were similar when reconstituted in 2:1:1 POPC:POPE:POPG and 3:1 POPC/DPOCL liposomes (time constants of 2.1 ± 0.9 s and 2.3 ± 1.1 s, respectively) (Fig. 1C). In contrast, ELIC5 activation was ∼50x slower in POPC liposomes (time constant of 123 ± 28 s) (Fig. 1C). Peak agonist responses were ∼20x higher in 2:1:1 POPC:POPE:POPG and 3:1 POPC/DPOCL liposomes compared to POPC liposomes, and no evidence of desensitization was observed in the three lipid conditions (Fig. 1C). Activation rates of ELIC5 in POPC liposomes did not vary with changes in the peak agonist response when reconstituting different quantities of ELIC5 (Supplementary Fig. 2A). Therefore, the slow activation of ELIC5 in POPC liposomes is not a consequence of lower reconstitution efficiency.

Since ELIC5 does not desensitize, we next examined the effects of lipid composition on steady-state ELIC5 activity as a function of agonist concentration. Here, we also tested binary mixtures of 3:1 POPC:POPE and 3:1 POPC:POPG. ELIC5 activity at saturating agonist concentration was ∼20x higher in 2:1:1 POPC:POPE:POPG and 3:1 POPC:DPOCL compared to POPC liposomes (Fig. 1D). 3:1 POPC:POPE also produced high ELIC5 activity, while 3:1 POPC:POPG produced intermediate activity (Fig. 1D). The agonist EC_50_ values (concentration of propylamine required for half-maximal activity) were similar in all lipid conditions, except 3:1 POPC:POPG which showed a shallow dependence that could not be fit using the Hill equation (Supplementary Figure 2B). The steepness of the dose response curves was higher in liposomes that fully support ELIC5 activity (2:1:1 POPC:POPE:POPG, 3:1 POPC:DPOCL, 3:1 POPC:POPE) compared to POPC or 3:1 POPC:POPG liposomes (Supplementary Figure 2B). Overall, the results indicate that PE and CL are sufficient to support ELIC5 activity.

We also tested whether ELIC5 activity can be rescued after reconstitution in POPC liposomes. ELIC5 in POPC liposomes was mixed with either empty POPC liposomes (control sample) or 2:1:1 POPC:POPE:POPG liposomes (test sample) and treated with five freeze-thaw cycles (Supplementary Figure 3A). This freeze-thaw treatment is expected to produce vesicle fusion leading to the introduction of some POPE and POPG into the test sample. The test sample exhibited significantly higher ELIC5 activity in response to agonist than the control sample (Supplementary Figure 3B), demonstrating that ELIC5 activity can be rescued after reconstitution in POPC liposomes and that the channel is not irreversibly inactivated.

### PE/PG and CL stabilize the resting and open-channel conformations of ELIC5

To investigate the structural basis by which PE/PG and CL support ELIC5 activity, we determined single-particle cryo-electron microscopy (cryo-EM) structures of ELIC5 reconstituted into spNW15 nanodiscs composed of 2:1:1 POPC:POPE:POPG, 3:1 POPC:DPOCL or POPC lipids. Structures were determined in the absence or presence of agonist (10 mM propylamine). ELIC5 in 2:1:1 POPC:POPE:POPG and 3:1 POPC:DPOCL nanodiscs produced single, consensus structures in the absence or presence of agonist that are consistent with previously-reported resting and open-channel conformations of ELIC5 (Supplementary Figure 4-5, Supplementary Table 1).

In contrast, single particle analysis of ELIC5 reconstituted in POPC nanodiscs showed increased structural heterogeneity both in the absence and presence of agonist. In the absence of agonist, three-dimensional variability analysis (3DVA) demonstrated continuous conformational heterogeneity, with motions resembling agonist-dependent changes including movement of Loop C, which caps the agonist binding site, and the TMD helices (Fig. 2A, Supplementary Video 1 and 2, Supplementary Figure 6). Based on the first component from the 3DVA, which showed the greatest variability in protein conformation (Fig. 2A), we partitioned the dataset into two classes to generate refined structures. Non-uniform refinement of these classes produced two structures: “class 1 unliganded” which is like the resting structure of ELIC5, and “class 2 unliganded” which is distinct with conformational changes resembling agonist-bound structures (Supplementary Figure 4-5). In “class 2 unliganded”, Loop C shows an inward capping, while the interfacial loops (β8-β9, β1-β2), M2-M3 linker, and TMD helices show conformations intermediate between the resting and open-channel structures (Fig. 2B-2F). Notably, the pore-lining M2 helix in “class 2 unliganded” is partially tilted with an intermediate conformation between the resting and open-channel structure. The pore-facing 9’ leucine (L240), which forms the activation gate in ELIC, orients towards the pore unlike the open-channel structure where L240 orients towards the adjacent subunit (Fig. 2F). Accordingly, the pore at 9’ has a radius of 2.5 Å, which is less constricted than the resting structure (1.9 Å) but more constricted than the open-channel structure (5 Å) (Supplementary Figure 7A). Since the 3DVA shows a continuum of conformations, it is likely that the particles in “class 2 unliganded” have significant conformational heterogeneity, which we cannot resolve by 3D classification either with C5 or C1 symmetry. Given the heterogeneity of this dataset and the difference in pore structure between “class 2 unliganded” and the open-channel structure, we suspect that the conformations sampled in the unliganded dataset are non-conducting. In summary, the resting state of ELIC5 is destabilized in a POPC lipid environment. The ion channel visits a continuum of states with structural features characteristic of partial activation yet not reaching the fully activated, open-channel state. We suggest that these states are less responsive to agonist consistent with the decreased activity of ELIC5 in POPC liposomes.

**Figure 2:**
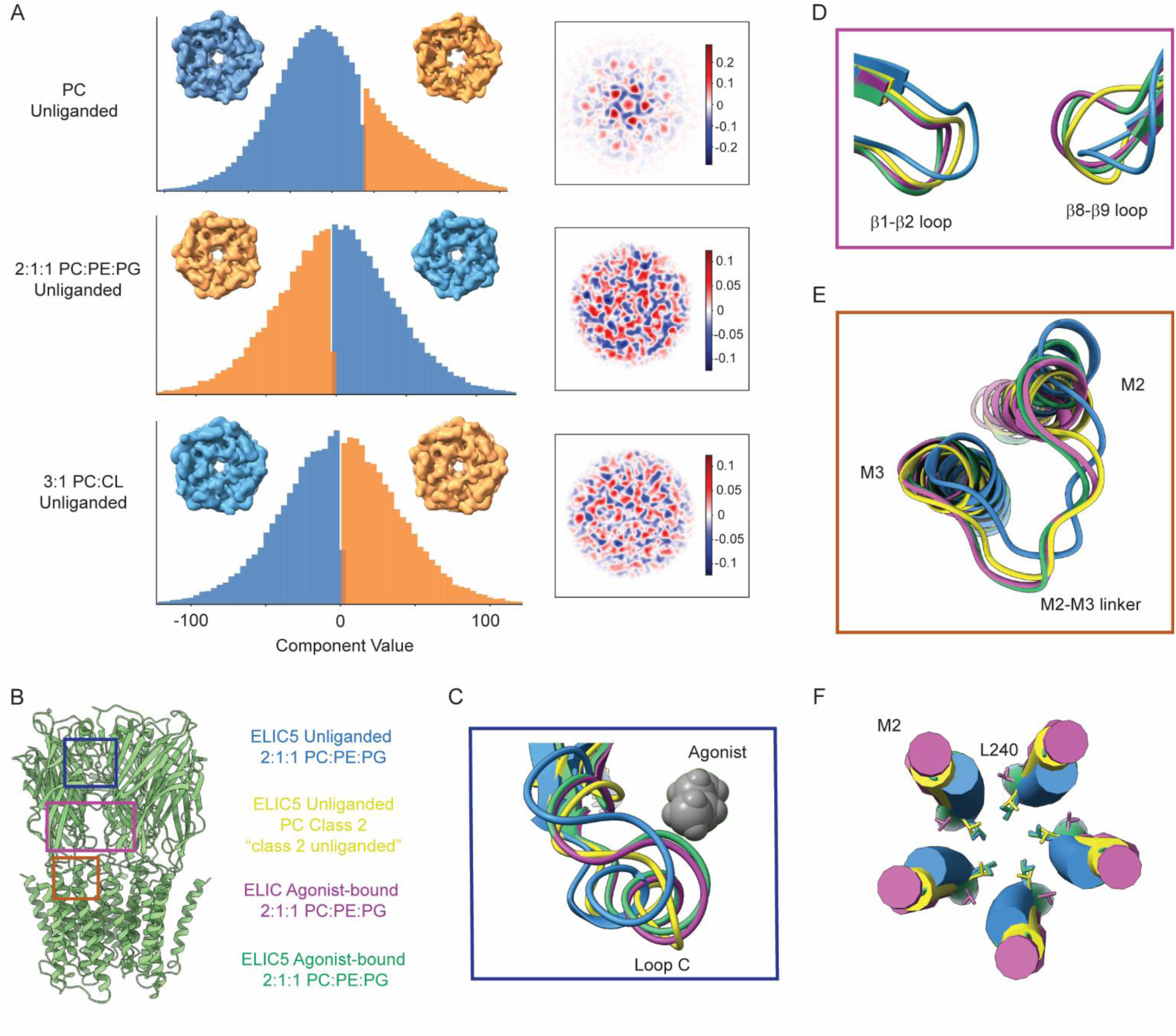
Structural effects of POPE/POPG and DPOCL on the unliganded structure of ELIC5. (A) *Left:* Histograms showing the distribution of particles along the first variability component (i.e. reaction coordinates) from 3DVA for ELIC5 in a POPC, 2:1:1 POPC:POPE:POPG and 3:1 POPC:DPOCL lipid environment. To illustrate differences between the datasets, the variability is separated into two clusters using cluster mode in cryoSPARC, and the 3D reconstruction for each cluster is shown with a view down the pore axis. *Right:* Representation of the first variability component of a central slice of ELIC5 from each dataset. Red and blue indicate positive and negative values of variability. (B) Structure of ELIC5 with boxes over the following regions being displayed: (C) Loop C which caps the agonist binding site, (D) the β1-β2 and β8-β9 loops in the ECD that interface with the TMD, and (E) the M2-M3 linker, and M2 and M3 helices. The structures being compared are: unliganded ELIC5 in 2:1:1 POPC:POPE:POPG (this study), unliganded ELIC5 Class 2 in POPC (this study, “class 2 unliganded”), agonist-bound WT ELIC in 2:1:1 POPC:POPE:POPG (PDB 8F34), and agonist-bound ELIC5 in 2:1:1 POPC:POPE:POPG (this study). (F) View of the M2 helices along the pore axis showing the side chain of L240 (9’).

In the presence of agonist, single particle analysis of ELIC5 in POPC nanodiscs also showed significant structural heterogeneity. After 3DVA and 3D classification, only two classes produced high resolution structures by non-uniform refinement (class 1 + agonist, class 2 + agonist). “Class 1 + agonist” is like the open-channel structure of ELIC5, and “class 2 + agonist” shows minor differences in the TMD compared to the open-channel structure. In “class 2 + agonist”, there is a reorientation of L240 (9’) producing a pore radius of 4.2 Å, which is narrower than the open-channel structure (Supplementary Figure 7B and 8B). Like the unliganded structures, the pore dimensions of this structure should be interpreted with caution because of the structural heterogeneity in the dataset. Notably, M4 is not resolved in the agonist-bound structures of ELIC5 in POPC (Supplementary Figure 8A). Thus, agonist-bound ELIC5 in POPC shows greater structural heterogeneity than in 2:1:1 POPC:POPE:POPG or 3:1 POPC:DPOCL, which likely relates to the low activity in a POPC environment. A definitive non-conducting structure of agonist-bound ELIC5 was not captured in the POPC dataset likely due to limitations in resolving the structural heterogeneity.

### Role of an outer leaflet M3-M4 site in lipid modulation

In the unliganded and agonist-bound ELIC5 structures in 2:1:1 POPC:POPE:POPG and 3:1 POPC:DPOCL, lipid-like densities were observed in the outer TMD in a region spanning the M4, M3 and M1 helices (Supplementary Figure 9). These were not as apparent in the structures in POPC nanodiscs (Supplementary Figure 9). In the 3:1 POPC:DPOCL structures (unliganded and agonist-bound), there are 4 interconnected lipid-like densities, which likely represent part of a bound DPOCL molecule, and this is most prominent in the unliganded structure (Fig. 3A) (16). The strongest lipid density is in a hydrophobic groove between M3 and M4. Therefore, we hypothesized that occupancy of this M3-M4 groove by DPOCL or POPE supports ELIC5 agonist response.

**Figure 3:**
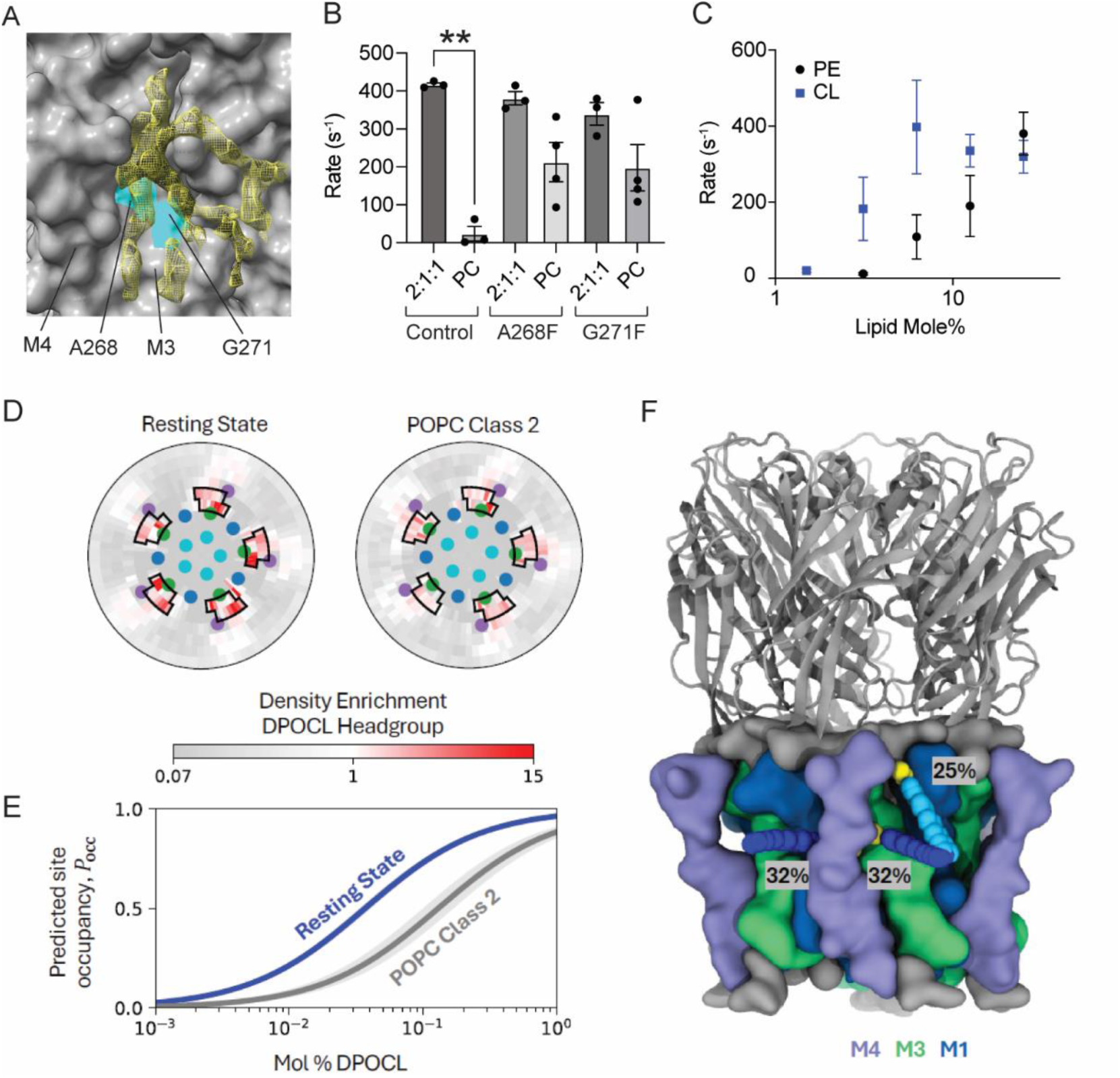
State-dependent binding of DPOCL to an outer leaflet M3-M4 site. (A) Surface representation of the unliganded ELIC5 structure in an 3:1 POPC:DPOCL lipid environment showing the cryo-EM density corresponding to a cardiolipin-like density with occupancy of an M3-M4 groove. The amino acids A268 and G271 are highlighted in this groove (cyan). (B) Tl^+^ flux rates of ELIC5 (control), ELIC5 + A268F, and ELIC5 + G271F in 2:1:1 POPC:POPE:POPG and POPC liposomes. Measurements were taken after a 20 min pre-incubation with 10 mM propylamine (n = 3-4). ** indicates p< 0.01. (C) Tl^+^ flux rates of ELIC5 in liposomes with varying POPE or DPOCL on a POPC background. Measurements were taken after a 20 min pre-incubation with 10 mM propylamine (n = 3-4). All data are shown as mean ± SEM for (n) independent experiments. (D) Radial distribution plots from CGMD simulations, measuring density enrichment of DPOCL headgroups in the outer leaflet within 5 nm of the protein center. DPOCL enrichment is observed near M3-M4 outlined in black. The colored dots represent the center of mass of each TMD helix. (E) Predicted occupancy of the boxed region near M4 as a function of mol % DPOCL in the membrane. The shaded region represents the 95% confidence interval with n=20 for the resting state and n=15 for POPC Class 2, which is the “class 2 unliganded structure”. (F) Molecular image of the resting structure of ELIC5 from the CGMD simulation with the average positions of most occupied clusters of DPOCL when its headgroup is near the outer leaflet M3-M4 binding site. Average DPOCL headgroup positions for each cluster are shown as yellow VdW spheres; average tail positions are shown as blue VdW spheres. Percentages correspond to the relative frequency of each binding mode.

To begin testing the role of the M3-M4 groove in the lipid sensitivity of ELIC, we identified A268 and G271 as two small residues in M3 that form the core of this groove (Fig. 3A). To perturb this site, we made A268F and G271F mutations on an ELIC5 background, introducing a bulky aromatic side chain in the M3-M4 groove. Both mutations increased channel activity in response to agonist in POPC liposomes (Fig. 3B). In other words, the mutations rendered ELIC more agnostic to its lipid environment. This result implicates the M3-M4 groove in determining the lipid requirements of ELIC activity. The aromatic substitutions may directly alter interactions between M3 and M4 (17) or lipid interactions at this site.

### CL supports ELIC5 activity by binding to the outer leaflet M3-M4 site

Cryo-EM densities of lipid in an outer leaflet M3-M4 site along with the effect of M3 mutations suggest the possibility that binding of DPOCL or POPE to this site mediates their effect on ELIC activity. To test this possibility further, we employed CGMD simulations to examine DPOCL or POPE interactions at this site. To guide design of the simulations, we first quantified the mole% of DPOCL and POPE required to support ELIC5 activity using the thallium flux assay. The effect of DPOCL saturated at ∼6.25%, while the effect of POPE reached a maximum at 25% without evidence of saturation (Fig. 3C). Liposomes could not be formed at greater than 25% POPE, which precluded measurements of ELIC5 activity at higher POPE concentrations.

Based on these results, we performed CGMD simulations of agonist-bound ELIC5 in membranes with three lipid compositions: 1) 95% POPC and 5% DPOCL, 2) 50% POPC, 25% POPE, and 25% POPG, and 3) 75% POPC and 25% POPE (Supplementary Figure 10 and 11). The open-channel cryo-EM structure of ELIC5 in 2:1:1 POPC:POPE:POPG liposomes was used for the first simulations, since this is a recently-determined structure of ELIC5 in the same membrane environment as was used in the thallium flux assay (11). In 95% POPC and 5% DPOCL, DPOCL was significantly enriched around the ELIC5 TMD. In the outer leaflet, DPOCL was enriched at the M3-M4 site, and in the inner leaflet, DPOCL was enriched at an M1-M4 site (Supplementary Figure 10 and 11). The CGMD simulations in 50% POPC, 25% POPE, and 25% POPG showed enrichment of POPG at approximately the same sites as DPOCL (14). However, POPE was only modestly enriched at these sites. Therefore, the CGMD results of the open-channel ELIC5 structure indicate that DPOCL binds to outer leaflet M3-M4 site more strongly than POPE. A potential limitation to this result is that coarse-grained beads may not sufficiently model the small headgroup of POPE.

The unliganded ELIC5 cryo-EM structures indicate that DPOCL stabilizes the resting conformation of ELIC5. We hypothesized that DPOCL stabilizes the resting conformation of ELIC5 by binding to the outer leaflet M3-M4 site. To test this possibility, we performed CGMD of the resting state structure of unliganded ELIC5 in 2:1:1 POPC:POPE:POPG, and “class 2 unliganded” which is the distinct unliganded structure only found in POPC. In the resting state, DPOCL was clearly enriched in the outer leaflet M3-M4 groove (Fig. 3D, Supplementary Figure 12). Clustering based on the DPOCL headgroup showed that DPOCL most frequently adopts one of three poses (Fig. 3F): the headgroup is buried behind M4 with the lipid tails in M3-M4 or M1-M4 grooves, or the lipid tails bind to the M3-M4 groove with headgroup interaction at the top of M4, M3 and M1. The latter is closest to the cryo-EM density of DPOCL (Fig. 3A). Strikingly, DPOCL shows greater enrichment at all three of these binding poses near M3-M4 in the resting state compared to the “class 2 unliganded” structure (Fig. 3D), with a density-threshold affinity (Δ𝐺_bind_) (18, 19) of -2.1 ± 0.1 kcal/mol for the resting state and -1.5 ± 0.3 kcal/mol for “class 2 unliganded”. These results are maintained when defining site boundaries that include a subset of poses (Supplementary Figure 12). The Δ𝐺_bind_ was used to predict the probability of site occupancy (𝑃_occ_) as a function of mole% DPOCL, demonstrating that DPOCL occupancy favors the resting state at all concentrations of DPOCL (Fig. 3E). We also observe enrichment of DPOCL at the inner leaflet M1-M4 site that favors the resting state structure over the “class 2 unliganded” structure (Supplementary Figure 13). These results indicate that DPOCL interaction with the outer leaflet M3-M4 and the inner leaflet M1-M4 sites are state-dependent, favoring the resting state structure over the distinct unliganded structure observed in POPC.

The CGMD results show enrichment of DPOCL at both outer and inner leaflet sites. To further test the possibility that DPOCL supports ELIC5 activity by binding to an outer leaflet site, we conducted a series of thallium flux experiments using symmetric and asymmetric liposomes. The cryo-EM structures and CGMD results indicate that DPOCL and POPG bind to overlapping sites in the outer leaflet localized to M3 and M4. Since POPG does not fully support ELIC5 activity, POPG may antagonize the effect of DPOCL presumably through competitive binding. Therefore, we examined ELIC5 activity in liposomes with 12.5% DPOCL and increasing %POPG. Indeed, ELIC5 activity decreased with increasing %POPG (Fig. 4A), suggestive of competitive displacement of DPOCL by POPG. In contrast, 25% POPG did not decrease ELIC5 activity in the presence of 25% POPE (Fig. 4A). While we cannot rule out the possibility that POPG antagonizes the effect of DPOCL through a non-competitive mechanism, the finding that POPG does not antagonize the effect of POPE argues against a non-competitive mechanism.

**Figure 4:**
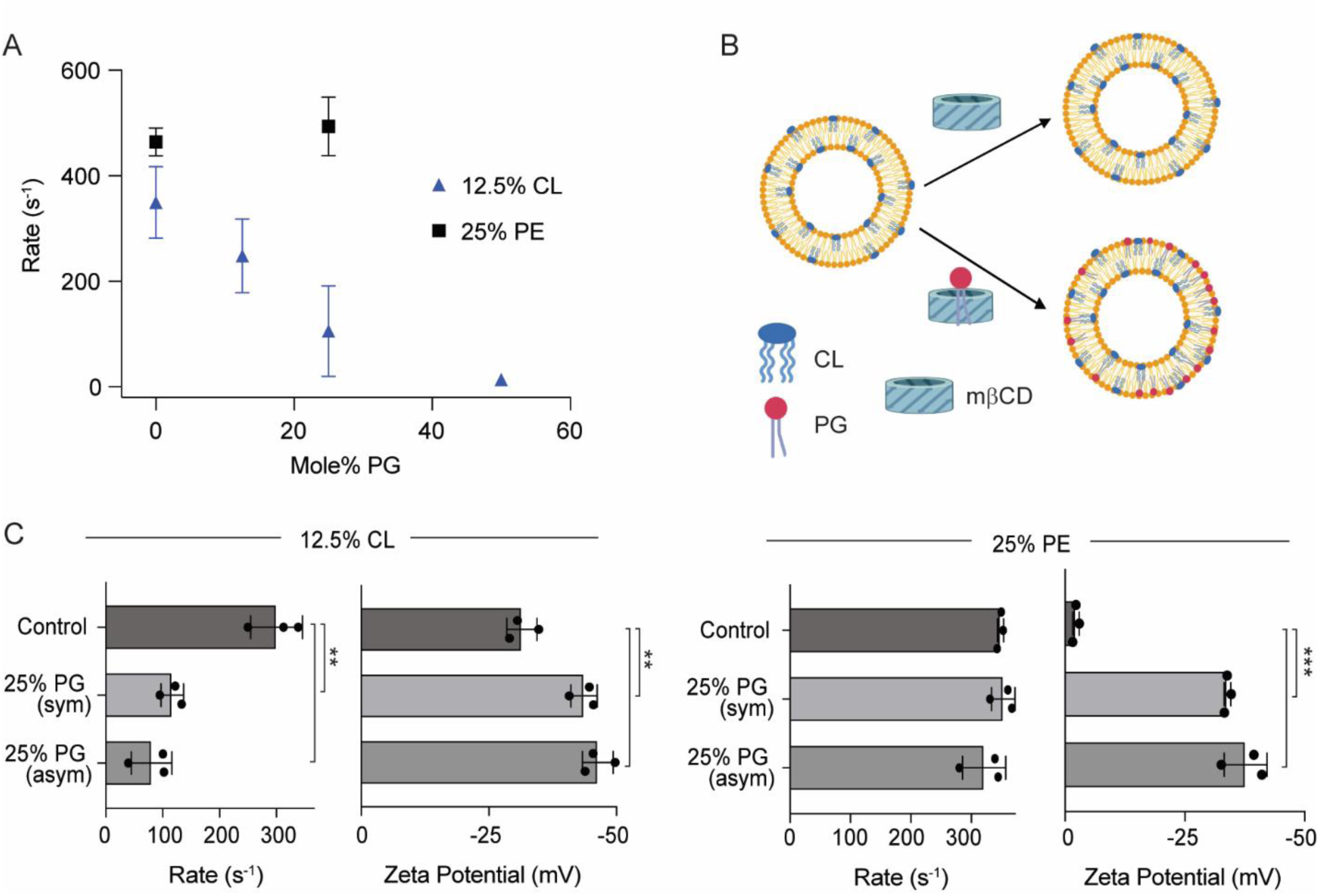
Outer leaflet POPG antagonizes DPOCL effects on ELIC5 activity. (A) Tl^+^ flux rates for ELIC5 in liposomes composed of 12.5% DPOCL or 25% POPE, and increasing mole% of POPG with a corresponding decrease in the mole% of POPC. Measurements were taken after a 20 min pre-incubation with 10 mM propylamine (n = 3). (B) Schematic of the asymmetric liposome experiment. ELIC5 proteoliposomes composed of 12.5% DPOCL or 25% POPE on a POPC background were treated with mβCD alone or mβCD loaded with POPG. The sample treated with mβCD loaded with POPG is anticipated to introduce 25% POPG to the outer leaflet in exchange for POPC. (C) *Left:* Tl^+^ flux rates and the corresponding zeta potentials of ELIC5 in liposomes with 12.5% DOPCL and no additional POPG (control), 25% POPG in both leaflets (sym), and 25% POPG in the outer leaflet (asym). *Right:* Same as *left* except all liposomes contain 25% POPE instead of 12.5% DPOCL. Measurements were taken after a 20 min pre-incubation with 10 mM propylamine (n = 3). All data are shown as mean ± SEM for (n) independent experiments. ** p<0.005 and *** p<0.0001 by one-way ANOVA with post-hoc Tukey comparison.

To test whether outer leaflet POPG can antagonize the effect of DPOCL in ELIC5, we reconstituted ELIC5 in liposomes with 12.5% DPOCL, and introduced 25% POPG to the outer leaflet using methyl-β-cyclodextrin-mediated lipid exchange (Fig. 4B) (13, 20, 21). Zeta potential measurements of the asymmetric proteoliposomes compared to control symmetric liposomes confirmed the introduction of 25% POPG to the outer leaflet (Fig. 4C). 25% outer leaflet POPG decreased ELIC5 activity like the result in symmetric liposomes (Fig. 4C). If indeed POPG is competitively displacing bound DPOCL, this finding indicates that the effect of DPOCL is mediated by an outer leaflet binding site. An equivalent experiment was performed with POPE where ELIC5 was reconstituted in liposomes with 25% POPE, and the effect of 25% outer leaflet POPG was examined. As was the case in symmetric liposomes, 25% outer leaflet POPG did not antagonize the effect of POPE (Fig. 4C).

## Discussion

We examined the mechanism by which PE and CL support ELIC activity using a combination of structural, computational and functional approaches, and taking advantage of a non-desensitizing ELIC mutant, ELIC5. ELIC5 allowed us to study the effect of phospholipids on agonist response without the complicating effect of desensitization. The cryo-EM structures indicate that POPE/POPG and DPOCL stabilize singular resting and open-channel conformations of ELIC5, while the structures in a POPC environment have increased structural heterogeneity. Most notably, the 3D variability analyses of unliganded ELIC5 show that the channel visits a continuum of states with structural features characteristic of agonist-induced changes. The “class 2 unliganded” structure of ELIC5 in POPC is not the same as the desensitized structure of ELIC (indeed, ELIC5 does not desensitize), but it does bear some resemblance. Therefore, it is reasonable to expect that these variable unliganded states of ELIC5 in POPC will render the channel less responsive to agonist, like a desensitized state.

In the unliganded and agonist-bound structures of ELIC5 in 3:1 POPC:DPOCL nanodiscs, we observed lipid densities consistent with bound DPOCL in an outer leaflet site adjacent to M3 and M4. Considering that: 1) mutations in this site alter ELIC5 lipid sensitivity, 2) DPOCL binding to this site favors the resting structure by CGMD, and 3) outer leaflet POPG antagonizes the effect of DPOCL, the results suggest that DPOCL supports ELIC5 activity by interacting with this outer leaflet site. Based on these findings, we propose a model for phospholipid modulation of ELIC activity (Fig. 5). In the presence of CL or PE, ELIC adopts a resting state that is responsive to agonist. In the absence of CL or PE, there is a destabilization of the resting state that decreases the efficacy of channel activation/opening. CL stabilizes the resting state of ELIC by binding to an outer leaflet M3-M4 site, which overlaps with the binding site(s) of PG. In addition, PG and CL decrease ELIC desensitization, stabilizing the activated state of the channel relative to the desensitized state, also by occupying this outer leaflet site (13). It is interesting to note that the outer leaflet M3-M4 site is a binding site for the polyunsaturated fatty acid, docosahexaenoic acid, which inhibits agonist response in ELIC (12). In contrast, PE supports ELIC activity by binding to an outer leaflet site not occupied by PG or an inner leaflet site, or by affecting the bulk properties of the membrane. CGMD simulations show that phospholipids interact strongly at an inner leaflet M1-M4 site in ELIC (22, 23) and the nAChR (24), and our results cannot rule out the possibility that inner leaflet phospholipids at this site or others also play a role in shaping agonist responses.

**Figure 5:**
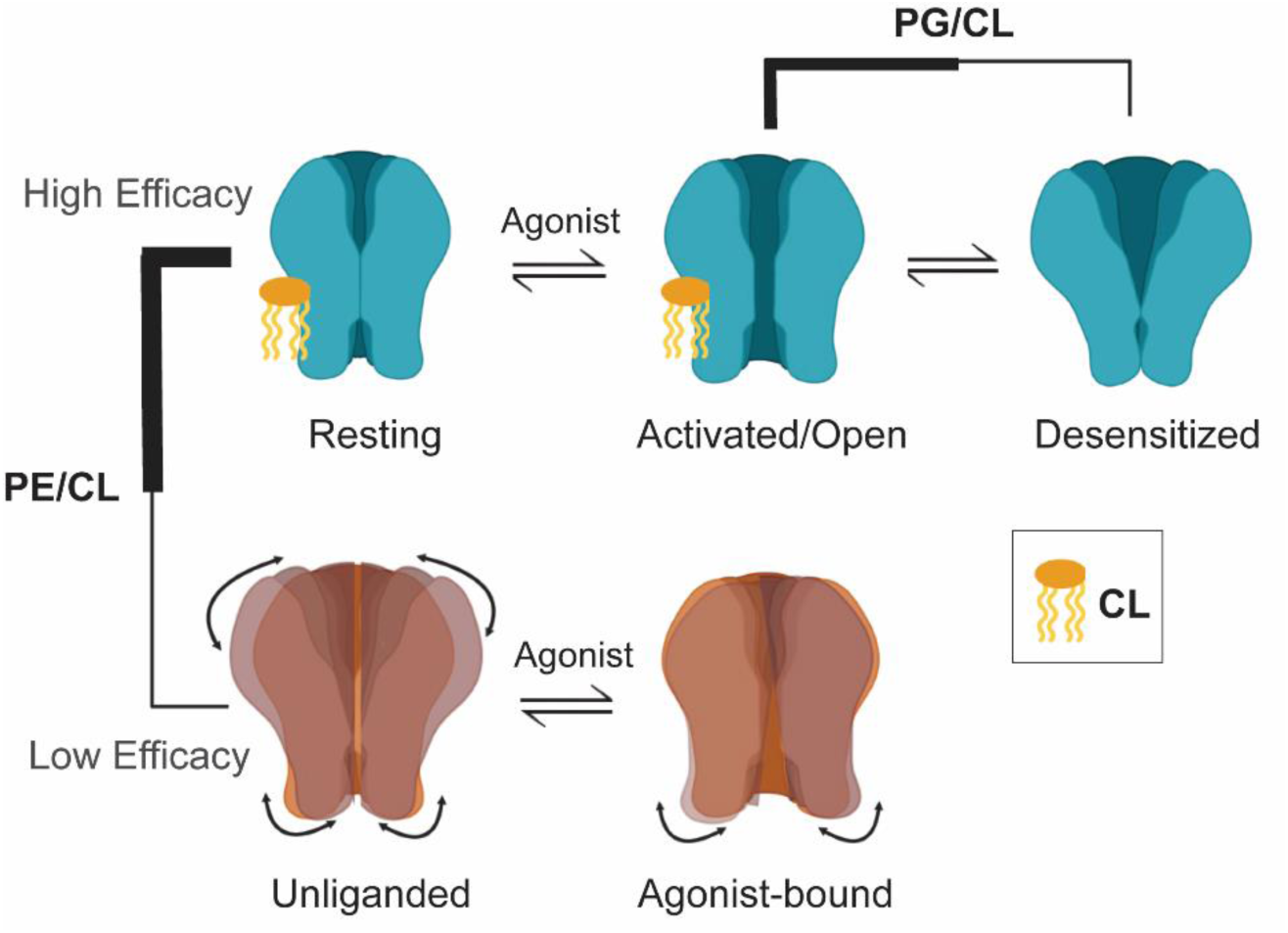
Model for phospholipid modulation of a pentameric ligand-gated ion channel. The data support a model where PE and CL stabilize the resting state of ELIC, promoting high efficacy ligand-activation. In the absence of PE and CL, the resting state is destabilized reducing the efficacy of ligand-activation. This is evidenced by the structural heterogeneity observed in the unliganded channel, which shows agonist-dependent changes that resemble a desensitized, unresponsive state. CL stabilizes the resting state by binding to an outer leaflet M3-M4 site. Binding of PG and CL to this site also decreases ELIC desensitization.

It has been estimated that <10% of nAChR at the neuronal cell membrane open when stimulated by agonist (25–27). This could be a result of receptors being distributed in distinct lipid environments (i.e. environments lacking PE, PS or cholesterol) that render the majority of the ion channels less responsive to agonist or functionally uncoupled (5, 6, 28). Other pLGICs such as the GABA_A_R and 5-HT3AR are dynamically distributed between cholesterol-rich lipid rafts and less-ordered lipid microdomains, and such differential lipid environments also appear to impact agonist responses in these ion channels (2, 4). The similar lipid sensitivity of ELIC and nAChR (i.e. both pLGICs show low agonist responses in PC membranes) (17) suggest a conserved mechanism of lipid modulation. Such a mechanism could be as we observe in ELIC, where the unliganded channel in a PC membrane adopts a continuum of conformations resembling but not equivalent to agonist-bound open or desensitized structures. Indeed, it was proposed that the unresponsive or uncoupled state of the nAChR in a PC membrane has some similarity to the desensitized state (29) yet is structurally distinct (30, 31). We conclude that, in the absence of agonist, ELIC is prone to visit many conformational states if the resting state is not stabilized by CL or PE/PG. This instability of the resting state in certain lipid environments may be a common phenomenon of pLGICs including the nAChR.

A notable feature of ELIC5 in PC membranes is the structure of M4. In the unliganded channel, M4 is dynamic moving towards the lipid membrane and away from the M1 and M3 helices. Similarly, in the agonist-bound channel, M4 is so flexible it is no longer resolved. The same finding (an unresolved M4) was observed for ELIC mutations that promote desensitization (28). However, in the present study, the increased heterogeneity of M4 is due to a lack of lipids such as CL and PE/PG. A long-standing hypothesis of lipid modulation of pLGICs, first proposed in the nAChR, is that certain lipids support agonist responses by stabilizing M4 (32). The cryo-EM structures from this study of ELIC5 in a PC environment provide the first direct structural evidence in support of this hypothesis. In a PC environment, M4 is more dynamic and its interactions with the rest of the channel are presumably weakened. It is possible that this lipid-dependent uncoupling of M4 is what destabilizes the resting and open-channel states of ELIC. Interactions between M4 and the M3/M1 helices as well as the ECD-TMD interface (e.g. β6-β7 loop) dictate ELIC and nAChR function (17, 31, 32). Immediately adjacent to this nexus of interactions is M4 interfacing with the lipid bilayer. Therefore, the results of this study lend credence to the model that M4 conformational dynamics in response to the lipid environment determines the agonist response of ELIC and potentially other pLGICs such as the nAChR (24, 33, 34).

In gram-negative bacteria where ELIC is expressed, PE, PG and CL are present at abundances expected to support ELIC activity. Both PG and CL are present in the outer leaflet (35), where they would modulate agonist responses in ELIC. Whether outer leaflet lipids modulate the function of mammalian pLGICs such as the nAChR remains to be established. In the plasma membrane of mammalian cells, pLGICs are embedded in an asymmetric lipid environment that includes PE and anionic phospholipids such as PS. However, both PE and PS are believed to exist mostly in the inner leaflet (36). If the activities of mammalian pLGICs are modulated by outer leaflet lipids as proposed for ELIC, it is more likely that lipids abundant in the outer leaflet such as cholesterol would be involved. Indeed, CGMD simulations of the nAChR in neuronal-like membranes show enrichment of cholesterol at the outer leaflet M3-M4 site (18, 37, 38).

## Methods

### ELIC mutagenesis, expression and purification

The pET-26-MBP-ELIC plasmid (originally from Raimund Dutzler, Addgene #39239)(39) served as the template for both wild-type (WT) and mutant ELIC constructs. Point mutations were introduced using Quikchange site-directed mutagenesis, with all resulting plasmids verified by Sanger sequencing (Genewiz).

Protein expression for both WT and mutant ELIC followed established protocols(40). OverExpress C43 (DE3) E. coli cells (Lucigen) were transformed and cultivated in Terrific Broth (Sigma), with protein expression induced by the addition of 0.1 mM IPTG for approximately 16 hours at 18 °C. Cells were collected by centrifugation, resuspended in Buffer A (20 mM Tris, pH 7.5, 100 mM NaCl) supplemented with an EDTA-free protease inhibitor cocktail (Roche), and disrupted under high pressure (∼15,000 psi) using an Avestin C5 homogenizer. Cellular debris was removed by ultracentrifugation to isolate membrane fractions, which were then resuspended in Buffer A and solubilized with 1% n-dodecyl-β-D-maltoside (DDM; Anatrace). Solubilized membranes were batch-bound to amylose resin (New England Biolabs) for 2 hours. The resin was washed extensively with Buffer A containing 0.02% DDM and 0.05 mM TCEP (ThermoFisher Scientific) before eluting the protein with Buffer A supplemented with 40 mM maltose (Sigma). The MBP fusion tag was cleaved by overnight incubation at 4 °C with HRV-3C protease (ThermoFisher Scientific) at a ratio of 10 units per mg of ELIC. The digested proteins were subjected to size exclusion chromatography on a Sephadex 200 Increase 10/300 column (GE Healthcare Life Sciences), equilibrated with Buffer A containing 0.02% DDM.

### Nanodisc Reconstitution

Lipid mixtures of POPC, POPE, POPG and DPOCL (Avanti Polar Lipids) was prepared in chloroform, dried thoroughly in a desiccator overnight, then rehydrated in Nanodisc buffer A (10 mM HEPES, pH 7.5, 100 mM NaCl) to achieve a lipid concentration of 7.5 mg/ml. The lipid suspension was subjected to five cycles of freeze-thaw and extruded through 400 nm polycarbonate filters (Avanti Polar Lipids). Destabilization of the liposomes was achieved by the addition of DDM to a final concentration of 0.4%, and incubated at room temperature for 3 hours (13). ELIC5 and the corresponding nanodisc scaffold protein were then added at a molar ratio of 1:2:360 for ELIC5:spNW15:lipid (ELIC5 mutant). This assembly was adjusted to a final DDM concentration of 0.2% and incubated at room temperature for 90 minutes. To remove detergent, Bio-Beads SM-2 (Bio-Rad) were added and the mixture was nutated overnight at 4 °C. Assembled nanodiscs containing ELIC5 were further purified by size exclusion chromatography (Sephadex 200 Increase 10/300 column) equilibrated in Nanodisc buffer A. Prior to cryo-EM studies, samples were supplemented with propylamine to a final concentration of 10 mM and concentrated to 0.6–1.2 mg/ml. spNW15 scaffolds were kindly provided by Huan Bao (Addgene plasmids #173482 and #173483) and purified by Ni-NTA affinity chromatography as previously described(14) without an additional size exclusion step.

### Cryo-EM sample preparation and imaging

3 µl of ELIC5 reconstituted in spNW15 nanodiscs was applied to freshly plasma-cleaned Quantifoil R2/2 copper grids. The grids were treated with a hydrogen/oxygen plasma for 60 seconds using a Gatan Solaris 950 system immediately prior to sample application. Following sample deposition, grids were blotted for 2 seconds at 100% relative humidity before being plunge-frozen in liquid ethane with a Vitrobot Mark IV (Thermo Fisher Scientific). Imaging was primarily performed on a Glacios microscope at 200 kV, also equipped with a Falcon 4 detector. Data acquisition utilized the EPU software (versions 2.12.1.2782 or 3.1.0.4506REL) in counting mode. Movies were collected at a pixel size of 1.184 Å, with defocus values ranging from −1 to −2.4 µm. 45-frame movies were collected, corresponding to total doses of 50.09 electrons/Å².

Single particle cryo-EM datasets were processed in CryoSPARC 4.5.1(41). The general workflow included patch motion and CTF correction, particle selection initially using 2D templates followed by Topaz particle picking and several rounds of 2D classification (42). Ab initio models served as the starting point for multiple rounds of heterogeneous refinement (28), and the final reconstructions were produced using non-uniform refinement with C5 symmetry.

To examine conformational heterogeneity, finalized particle sets were symmetry expanded by 5-fold symmetry and subjected to 3D Variability Analysis (3DVA), employing a filter resolution of 6 Å and five principal component (PCA) modes. This was performed for all datasets, but only the datasets of unliganded and agonist-bound ELIC5 in POPC nanodiscs showed clear heterogeneity. For unliganded ELIC5 in POPC nanodiscs, the symmetry-expanded particles were analyzed using cluster mode in 3DVA, focusing on the primary mode of variability (43). This process yielded three distinct particle clusters, representing the major classes within this mode. To ensure uniqueness and data integrity, duplicate particles were identified and removed from each cluster. The resulting, non-redundant particle sets from each cluster were then independently subjected to non-uniform refinement for each cluster/class. Two of these classes were indistinguishable and therefore combined yielding the final structures of “class 1 unliganded” and “class 2 unliganded”.

Model building was performed by fitting an initial ELIC structure (PDB 8D66) into the cryo-EM density map using PHENIX 1.19.2 for real-space refinement, followed by manual building in COOT 0.9.6. Multiple rounds of real-space refinement and interactive corrections were performed iteratively in PHENIX and COOT. The agonist propylamine was modeled into the cryo-EM map and oriented based on its predicted binding mode (44).

### Assessment of Protein Orientation in Liposomes

The orientation of liposome reconstituted ELIC (A6C/C300S/C313S) was assessed using a fluorescence quenching assay (15). ELIC (A6C/C300S/C313S) was incubated with the thiol-reactive fluorophore DY-647P1-maleimide 10-fold of protein concentration overnight at 4 degrees C (Dyomics GmbH, Cat. No. 647P1-03). After labeling, free label was removed by size exclusion chromatography and the fluorescently-labeled protein was reconstituted into liposomes as described in the following section. Fluorescence measurements of the proteoliposomes were carried out on a Fluoromax spectrofluorometer. After recording baseline fluorescence, TCEP was added to a final concentration of 10 mM to quench fluorophores accessible on the external leaflet of the liposome membrane, reporting the fraction of ELIC with the ECD exposed to the extra-liposomal space (“outside-out” orientation). To determine the total population of labeled protein, Triton X-100 was subsequently introduced to permeabilize the liposomes, granting TCEP access to fluorescently-labeled ELIC with the “inside-in” orientation.

### Stopped-Flow Thallium Flux Assay

The stopped-flow thallium flux assay of ELIC proteoliposomes were conducted using a sequential-mixing protocol with an SX20 stopped-flow spectrofluorometer (Applied Photophysics) controlled by Pro-Data SX software, as previously described (23, 45). Liposomes were prepared by drying 7.5 mg of the desired lipid in glass vials to a thin film under nitrogen, then subjecting them to further desiccation under vacuum overnight. The following day, lipids were hydrated in 500 μl of reconstitution buffer (100 mM NaNO₃, 10 mM HEPES, pH 7.0; Buffer R) with the addition of 250 μl of 75 mM ANTS (8-Aminonaphthalene-1,3,6-trisulfonic acid; ThermoFisher Scientific) and ∼18 mg of CHAPS (Anatrace). This mixture was vortexed and sonicated with heat until fully solubilized. Once equilibrated to room temperature, ELIC protein (1 μg protein per mg lipid) was incorporated into the lipid mixture and incubated for 30 minutes at room temperature. 3 μg protein per mg lipid was used for the ELIC5 POPC freeze-thaw experiment, and for the fluorescently-labeled ELIC (A6C/C300S/C313S) experiment. Bio-Beads SM-2 (1 ml of 1:1 v/v slurry in Buffer R) and an additional 250 μl of 75 mM ANTS were added, and the mixture was incubated for 2.5 hours at room temperature with gentle mixing. The resulting proteoliposomes were extruded through a 0.1 μm polycarbonate membrane (Avanti) using a mini-extruder (20 passes) and then stored overnight at room temperature with gentle rotation. The following day, the liposomes were diluted to 2.5 ml in stopped-flow assay buffer (140 mM NaNO₃, 10 mM HEPES, pH 7.0; Stopped-flow Buffer A), loaded onto a PD-10 desalting column (Cytiva), eluted with 3 ml of Stopped-flow Buffer A, and further diluted five-fold in the same buffer. For flux assays, liposomes containing ELIC were rapidly mixed 1:1 with Stopped-flow Buffer A containing propylamine at twice the desired final concentration to initiate channel activation. After variable preincubation times (10 ms to 25 s), the sample was mixed 1:1 with quenching buffer (90 mM NaNO₃, 50 mM TlNO₃, 10 mM HEPES, pH 7.0), and ANTS fluorescence (excitation at 360 nm, emission >420 nm) was recorded for 1 s.

For ELIC5 flux measurements under pre-equilibration with agonist (propylamine), Proteoliposomes containing ELIC5 were incubated with agonist at the indicated final concentration for 20 minutes at room temperature prior to measurement. After this pre-incubation, samples were subjected to a single-mixing stopped-flow sequence using the SX20 fluorometer, wherein the agonist-pretreated proteoliposomes were rapidly mixed 1:1 with quenching buffer (90 mM NaNO₃, 50 mM TlNO₃, 10 mM HEPES, pH 7.0) supplemented with an equal concentration of agonist immediately before fluorescence acquisition. For each data point, measurements were repeated at least 3 times from independent preparations.

### Preparation and Analysis of Asymmetric Liposomes

Asymmetric liposomes were generated by modifying the standard liposome preparation protocol described in the Stopped-flow thallium flux assay section (13). For these experiments, the protocol was scaled up two-fold to ensure adequate sample for both the exchange and no-exchange conditions. After preparing a 2x larger preparation of ELIC5 proteoliposomes using Biobeads and storing the proteoliposomes overnight at room temperature, the total liposome preparation was divided into two equal 1.5 ml aliquots. To introduce POPG to the outer leaflet of these proteoliposomes, the required concentrations and volumes of donor POPG and methyl-β-cyclodextrin (MβCD, Sigma) were determined using spreadsheets provided by Markones et al (21). The calculations were based on a post-extrusion lipid concentration of 6.66 mM in 1.5 ml acceptor liposomes. For the POPG transfer, a MβCD-POPG complex (25% MβCD saturation with POPG) was prepared by combining 167.5 µl of 7.5 mM POPG in stopped-flow buffer, 650 µl of 300 mM MβCD, and 126 µl stopped-flow buffer. The mixture was incubated at 50 °C with shaking (1,000 rpm, Eppendorf thermomixer) for 20 min. In parallel, a control solution omitting POPG was assembled by substituting buffer in place of POPG. Both mixtures and the thermomixer were then cooled to 28 °C. The 1.5 ml aliquots of acceptor liposomes were each mixed with 1 ml of either MβCD-POPG (for exchange) or the POPG-free MβCD solution (for no-exchange control), followed by incubation at 28 °C, rotating at 400 rpm for 20 min. Each 2.5 ml sample was passed through a PD-10 column to remove extra-liposomal ANTS and then characterized in by the stopped-flow thallium flux assay as described above. Each replicate represents an independent preparation.

### Zeta Potential Measurement

ζ potential was employed to assess net surface charge and validate successful incorporation of POPG into the outer leaflet of the proteoliposomes, using methodologies adapted from Markones et al. ((13, 21)). Measurements were performed on a Malvern Zetasizer Nano-ZS ZEN 3600, equipped with a flow-through high concentration zeta potential cell (HCC), operating under Zetasizer nano software v3.30. The system was pre-equilibrated to 28 °C for 20 min prior to sample analysis. For each measurement, instrument settings included a 28 °C measurement temperature, a 90 s equilibration, and 10–100 runs per sample with water set as the dispersant. Immediately after passing the proteoliposomes through the PD-10 column, 100 µl of sample was diluted to 1 ml with 100 mM HEPES (pH 7); the final MβCD concentration was approximately 4.3 mM. The sample was loaded into a 1 ml syringe and injected into the HCC, ensuring displacement of air and old solution (with an additional ∼250 µl used for rinsing between runs). Each sample was measured in triplicate. The HCC cell was flushed with hot deionized water and dried under nitrogen before each new measurement and thoroughly cleaned post-measurement using 1% Hellmanex II in deionized water.

### CGMD Simulation Setup

We coarse-grained three structures of ELIC5 using the *Martinize* tool(46): agonist-bound ELIC5 in 2:1:1 POPC:POPE:POPG liposomes (PDB 9NGC), unliganded ELIC5 in 3:1 POPC:DPOCL nanodiscs (PDB 9YEN, this study), unliganded ELIC5 in POPC nanodiscs Class 2 (PDB 9YEW, this study). These structures putatively represent the activated/open-channel state, resting state, and an unliganded, low-efficacy state in POPC, respectively. We then embedded each structure in model membranes using *Insane* (47). The open-channel structure (PDB 9NGC) was embedded in two model membranes: one containing 2:1:1 POPC:POPE:POPG, and one containing 3:1 POPC:POPE (see Table 1). Each of the three structures were also embedded in 20:1 POPC:Cardiolipin (CL). To aid in the determination of binding site accessible area, each of the three structures were also embedded in 100% DPPC (19). To determine the ideal gas component of the binding affinity, “bulk” systems of each model membrane were also created without a protein embedded. These systems were all solvated with water and 15% NaCl. The initial box size was 25 x 25 x 20 nm^3^, allowing for the placement of approximately 1,000 lipids in each leaflet.

**Table 1:**
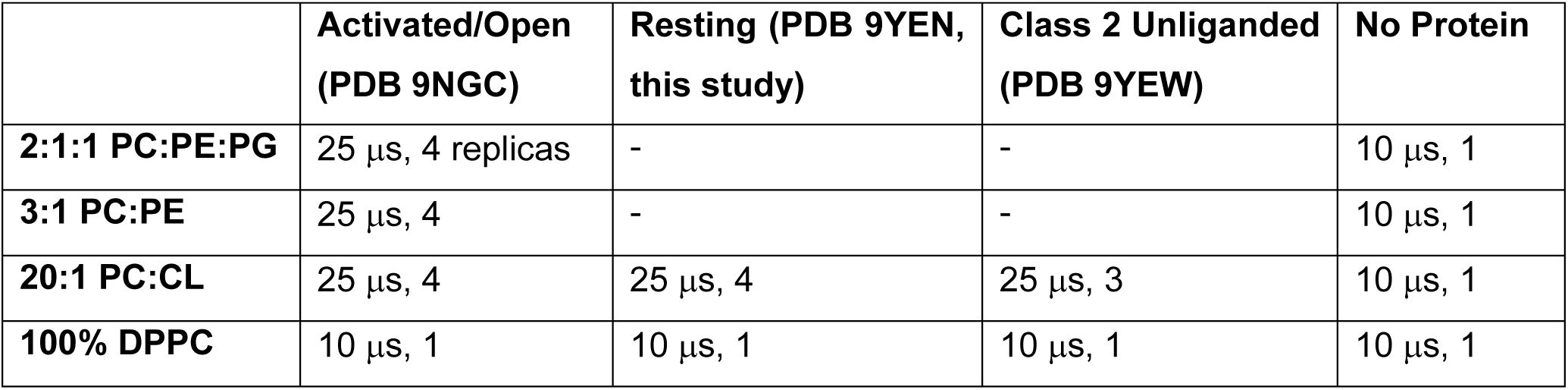
Description of simulation systems. Simulation length and number of replicas is specified for each protein structure simulated (columns) and each model membrane used (rows).

|  | **Activated/Open**<br>**(PDB 9NGC)** | **Resting (PDB 9YEN,**<br>**this study)** | **Class 2 Unliganded**<br>**(PDB 9YEW)** | **No Protein** |
| --- | --- | --- | --- | --- |
| **2:1:1 PC:PE:PG** | 25 μs, 4 replicas | - | - | 10 μs, 1 |
| **3:1 PC:PE** | 25 μs, 4 | - | - | 10 μs, 1 |
| **20:1 PC:CL** | 25 μs, 4 | 25 μs, 4 | 25 μs, 3 | 10 μs, 1 |
| **100% DPPC** | 10 μs, 1 | 10 μs, 1 | 10 μs, 1 | 10 μs, 1 |

The Density-Threshold Affinity (DTA) protocol detailed in Sandberg et. al (19) requires two additional simulations. Each structure was embedded in a 100% DPPC membrane and simulated for 10μs, one replica per structure. Each model membrane was also simulated without a protein embedded for 10 μs, one replica per model membrane.

All simulations were performed using GROMACS 2024 (48) and the martini 2.2 force field (46). The *v-rescale* thermostat was set to 320 K and a semi-isotropic *C-rescale* barostat was set to 1 bar in both directions. The compressibility modulus was 3×10^-4^ bar^-1^ with time constant 50 ps. Electrostatics used the *reaction-field* method and van der Waals were *cutoff* with a *Potential-shift-verlet* modifier, both with *r*-values of 1.1 nm. If a protein was present in the simulation, backbone beads were restrained to their initial positions with a force constant of 1,000 kJ mol^-1^ nm^-2^, and *refcoord_scaling* was set to *com* to avoid membrane compression (49). Systems underwent 10,000 steps of steepest-descent energy minimization, then 40,000 steps of MD with a time step of 1 fs, then MD with time step of 25 fs. The simulation length and number of replicas for each system are shown in Table 1.

### CGMD Simulation Analysis

Simulation trajectories were visualized in VMD(50) and further analysis was performed using TCL and python scripts (51). In all cases, the first 5 μs of the simulation trajectory was removed prior to analysis. As in previous studies (12, 18, 19), putative binding sites were identified by measuring the density enrichment of a lipid species (or a subsection of the lipid, e.g. its headgroup) within 5 nm of the protein center of mass. Density enrichments were measured on a polar lattice with 10 evenly spaced radial bins and 90 evenly spaced azimuthal bins. Contiguous lattice bins exhibiting high enrichment were selected as potential binding sites.

The Density-Threshold Affinity (ΔG_bind_) was then measured for each site, following a previously published protocol (19). The ΔG_bind_ was then used to calculate the probability that the site will be occupied (*P*_occ_) by some lipid as a function of its concentration,

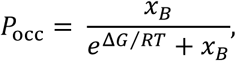

where 𝑥_𝐵_ is the mole% of the lipid in question. In the 20:1 POPC:DPOCL simulations, the ΔG_bind_ of DPOCL to identically defined sites around each of two protein structures/states is compared. To facilitate accurate comparisons across the two structures, the accessible area of each site in the resting state structure was used for analysis of both structures. Versions of these graphs without this correction are available in Supplementary Figure 12.

The most-occupied binding mode(s) were determined via cluster analysis. A custom TCL script was used to capture all instances across all replicas of a lipid species (or a specific component of that lipid species, e.g. its headgroup) entering the binding site and output them to one molecular trajectory. Each trajectory was then analyzed using the *measure cluster* command in VMD with number of clusters set to 8 and a cutoff of 6. In the case of DPOCL, the linker bead entering the site was the binding criterion and clustering was performed on the linker, phosphate, and glycerol bead positions. Clusters were then further refined by combining clusters that had linker beads within 4 Å of one another.

## Data Availability

The data supporting the findings of this study are available within the paper and supplementary information files. The cryo-EM maps have been deposited in the Electron Microscopy Data Bank (EMDB) under accession codes EMD-72856, EMD-72858, EMD-72859, EMD-72861, EMD-72865, EMD-72867, EMD-72869, and EMD-72875. The structural coordinates have been deposited in the RCSB Protein Data Bank (PDB) under the accession codes 9YEN, 9YEP, 9YEQ, 9YES, 9YEW, 9YEY, 9YF1, and 9YF3.

## Acknowledgements

This study was supported by NIH grant R35GM137957 to W.W.C., and a Foundation for Anesthesia Education and Research mentored training grant and Deans’ Scholars Fund to B.K.T.

## Authors Contributions Statement

B.K.T. and W.W.C. conceived the project and designed the experimental procedures. B.K.T. and H.X. carried out the protein expression and purification, nanodisc and liposome reconstitution and single particle analysis, and thallium flux assay. J.W.S. and G.B. carried out the coarse-grained molecular dynamics simulations and data analysis. All authors reviewed the paper.

## Competing Interests Statement

The authors declare no competing interests.

---

**Supplementary Figure 1:**
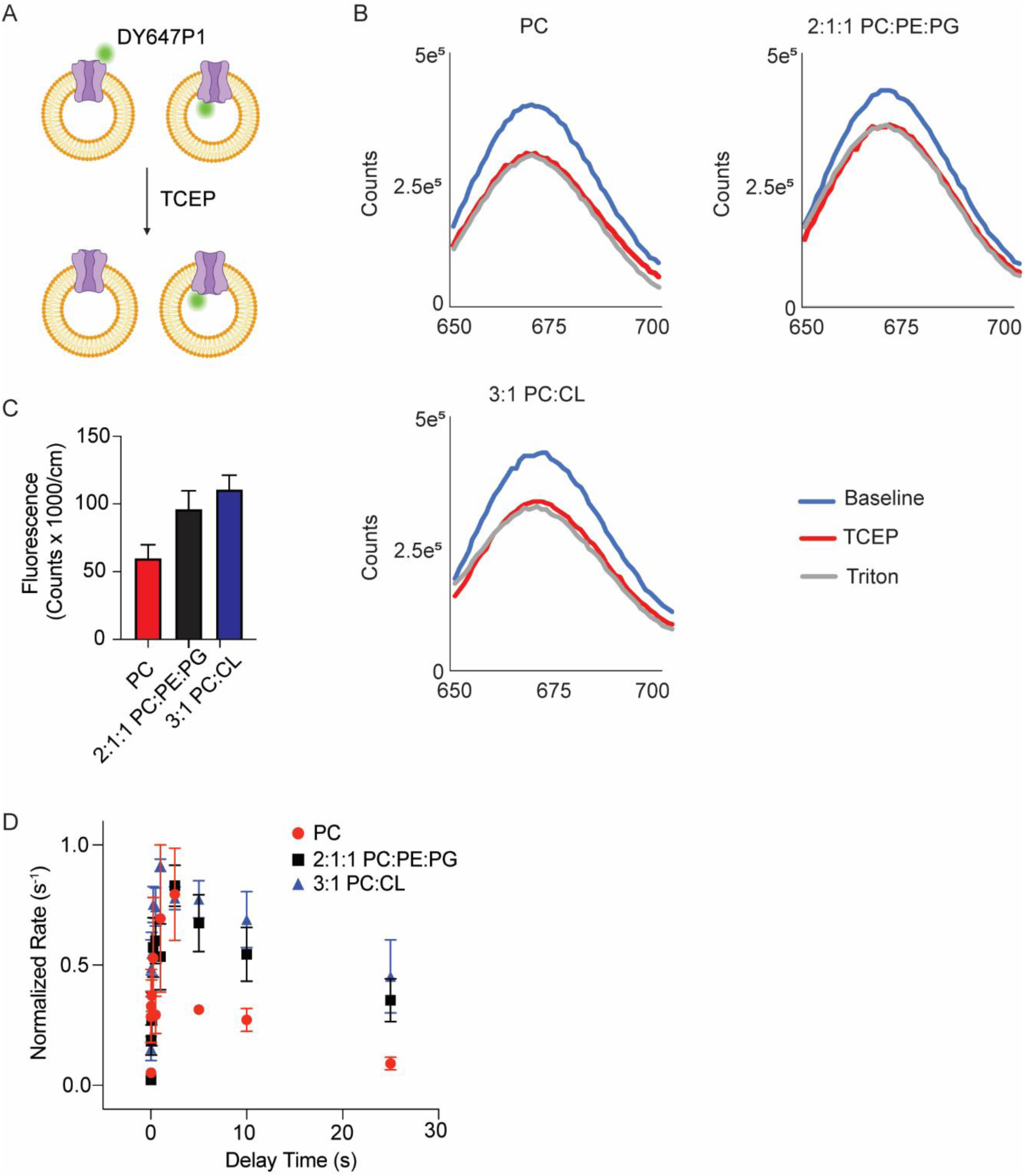
Fluorescence and Activity of DY647P1-labeled ELIC (A6C/C300S/C313S) in liposomes. (A) Schematic of ELIC (A6C/C300S/C313S) labeled with DY647P1 on the cysteine in position 6 and reconstituted in liposomes in both orientations. Treatment with TCEP only quenches those channels with the ECD facing the extra-liposomal space. (B) Fluorescence spectra of DY647P1-labeled ELIC (A6C/C300S/C313S) in liposomes with POPC, 2:1:1 POPC:POPE:POPG, and 3:1 POPC:DPOCL before TCEP, after TCEP, and after Triton X-100. The data indicate that most channels have the ECD facing the extra-liposomal space. The residual fluorescence after Triton X-100 treatment indicates that some fluorescently-labeled ELIC is not accessible to TCEP possibly because the protein is aggregated and not reconstituted in liposomes. To optimize the fluorescent signal, DY647P1-labeled ELIC (A6C/C300S/C313S) was reconstituted at 3 µg protein per 1 milligram lipid for this experiment. (C) The difference in fluorescence signal of DY647P1-labeled ELIC (A6C/C300S/C313S) before and after TCEP, indicating the quantity of ELIC with the ECD facing the extra-liposomal space (n = 3). (D) Tl^+^ flux rates of DY647P1-labeled ELIC (A6C/C300S/C313S) reconstituted in liposomes with POPC, 2:1:1 POPC:POPE:POPG, and 3:1 POPC:DPOCL. The data are normalized to the peak rate (n = 3-8, data from Figure 1B). All data are shown as mean ± SEM for (n) independent experiments.

**Supplementary Figure 2:**
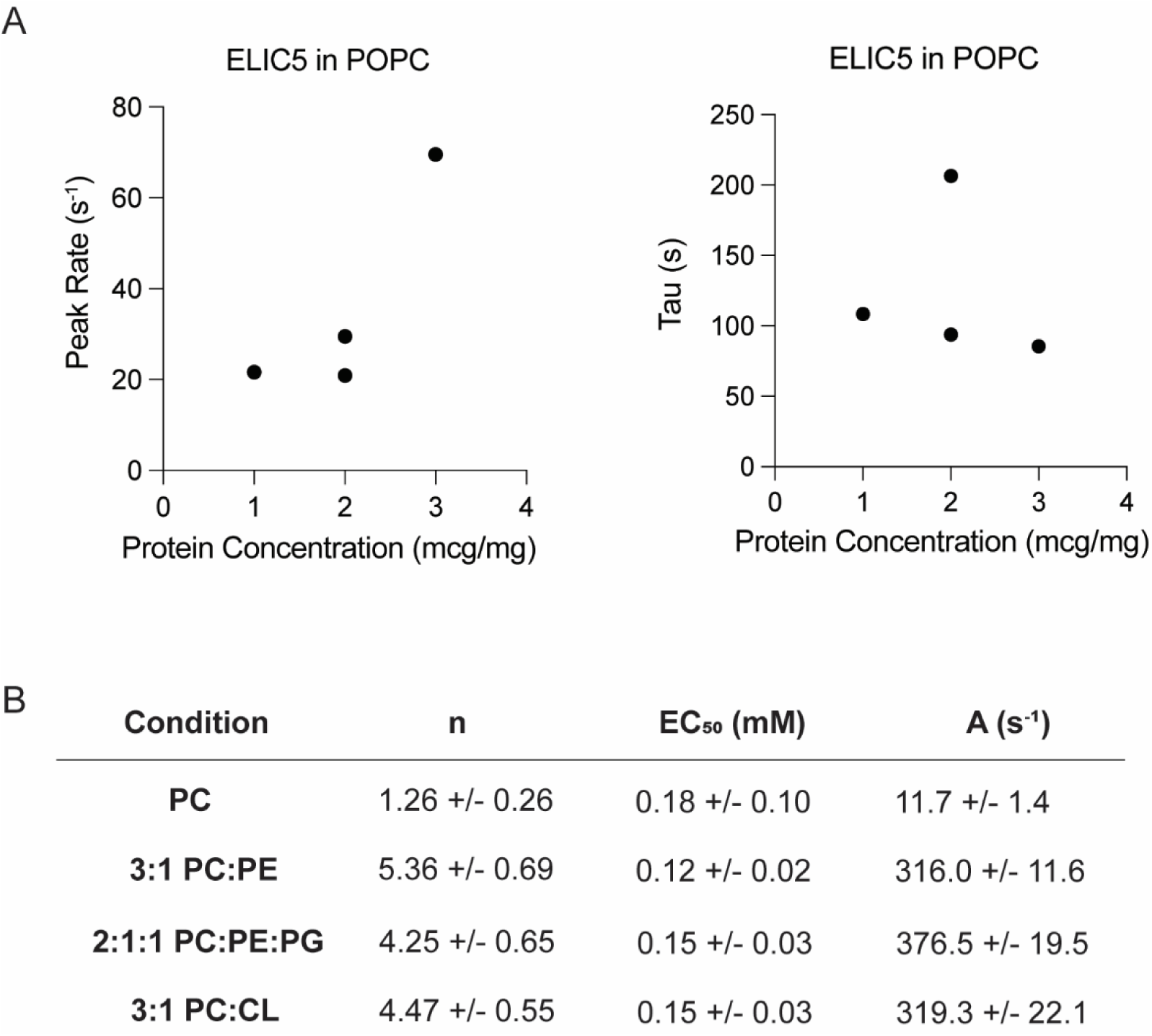
Relationship between ELIC5 peak agonist response or activation rate and the quantity of protein added to the liposome preparations. (A) To examine whether ELIC5 peak response or activation rate in POPC liposomes is dependent on the amount of protein used, ELIC5 activation was examined as in Figure 1C using different amounts of ELIC5 (micrograms of ELIC5 per milligram of lipid) for the liposome reconstitution. Shown are the peak Tl^+^ flux rates (*left*) and the time constants of activation (*right*) as a function of ELIC5 protein concentration. As expected, the peak agonist response increases with protein concentration but not the time constant of ELIC5 activation. (B) Results of fitting the Hill equation to the data in Figure 1D (n = 3-6, ± SD). n is the Hill coefficient, EC_50_ is the concentration of propylamine producing half maximal response, and A is the maximum activity (Tl^+^ flux rate).

**Supplementary Figure 3:**
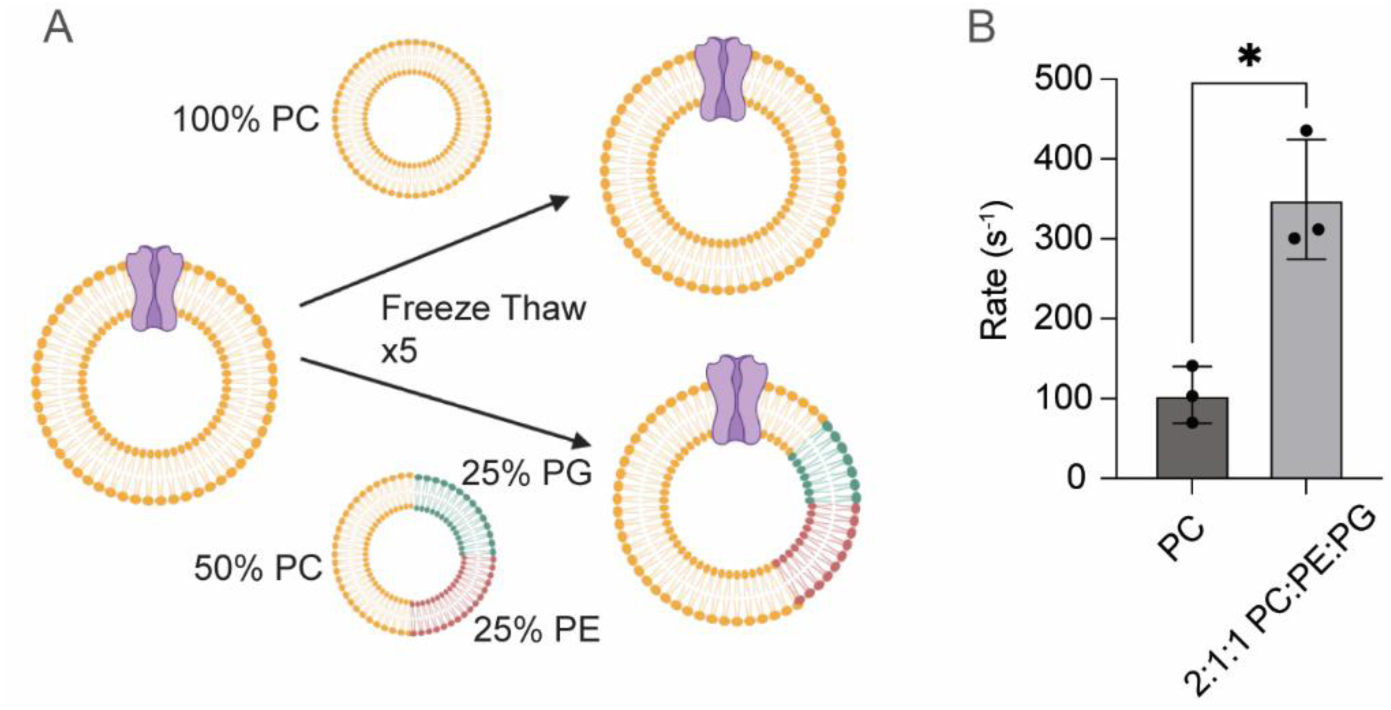
ELIC5 activity can be rescued after reconstitution in POPC liposomes. (A) Schematic of the experiment where ELIC5 is reconstituted in POPC liposomes. The preparation is split and mixed with equal quantities of POPC or 2:1:1 POPC:POPE:POPG liposomes. Each sample was then treated with 5 rounds of freeze-thaw. Due to liposome fusion, the anticipated result is ELIC5 proteliposomes with POPC-only or up to 12.5% POPE and 12.5% POPG. (B) Tl^+^ flux rates of ELIC5 from samples treated by freeze thaw with POPC liposomes or 2:1:1 POPC:POPE:POPG liposomes (n = 3). All data are shown as mean ± SEM for (n) independent experiments. * p<0.01 from paired T-test. To ensure adequate Tl^+^ flux rates, 3 micrograms of ELIC5 per milligram of lipid was used for each preparation. Unless otherwise specified, all other stopped-flow thallium flux experiments in this study used 1 microgram of ELIC5 per milligram of lipid.

**Supplementary Figure 4:**
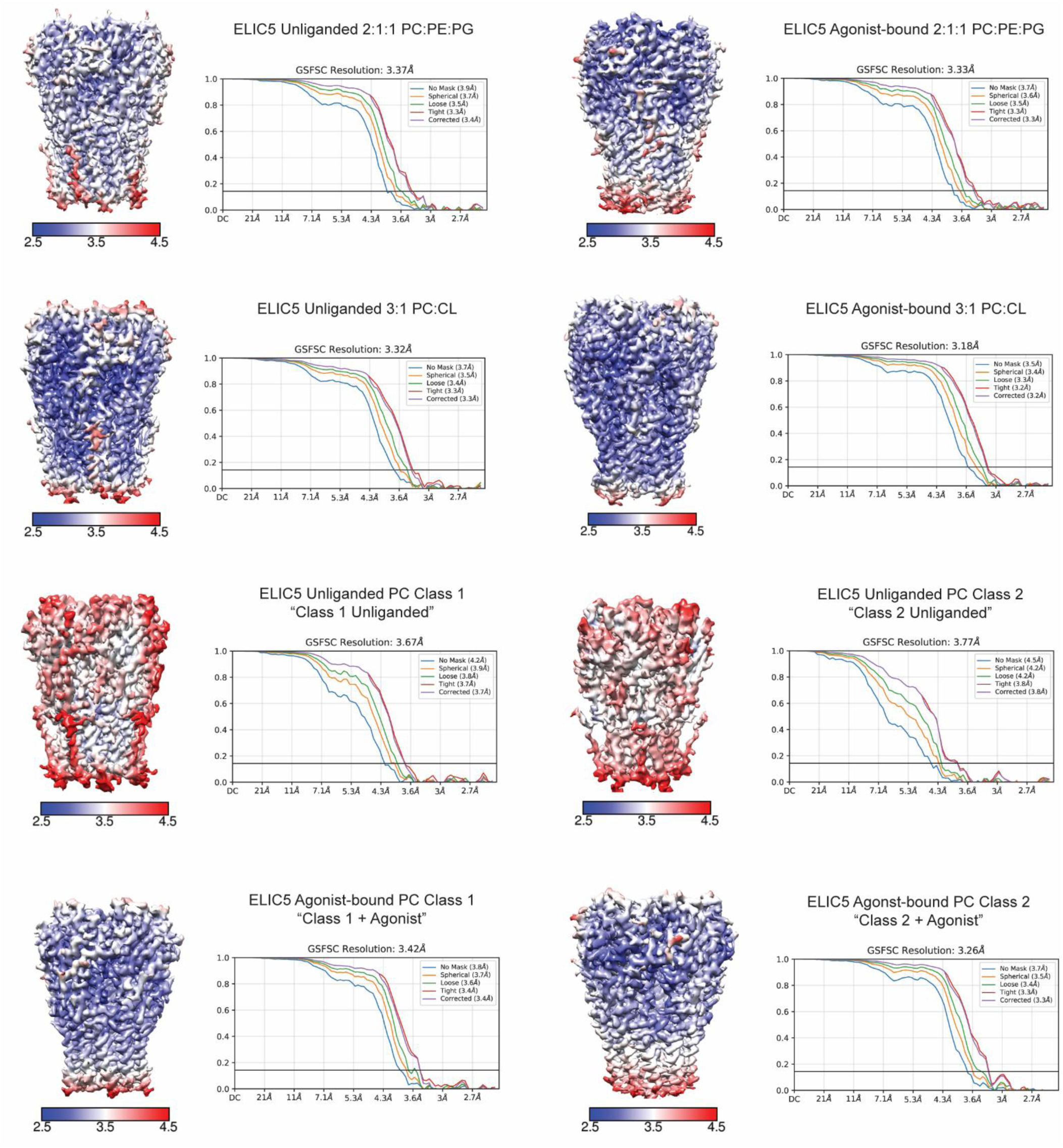
Local resolution and FSC curves for cryo-EM structures. For each structure, on the left is the cryo-EM map colored according to local resolution and on the right is the FSC curve from cryoSPARC.

**Supplementary Figure 5:**
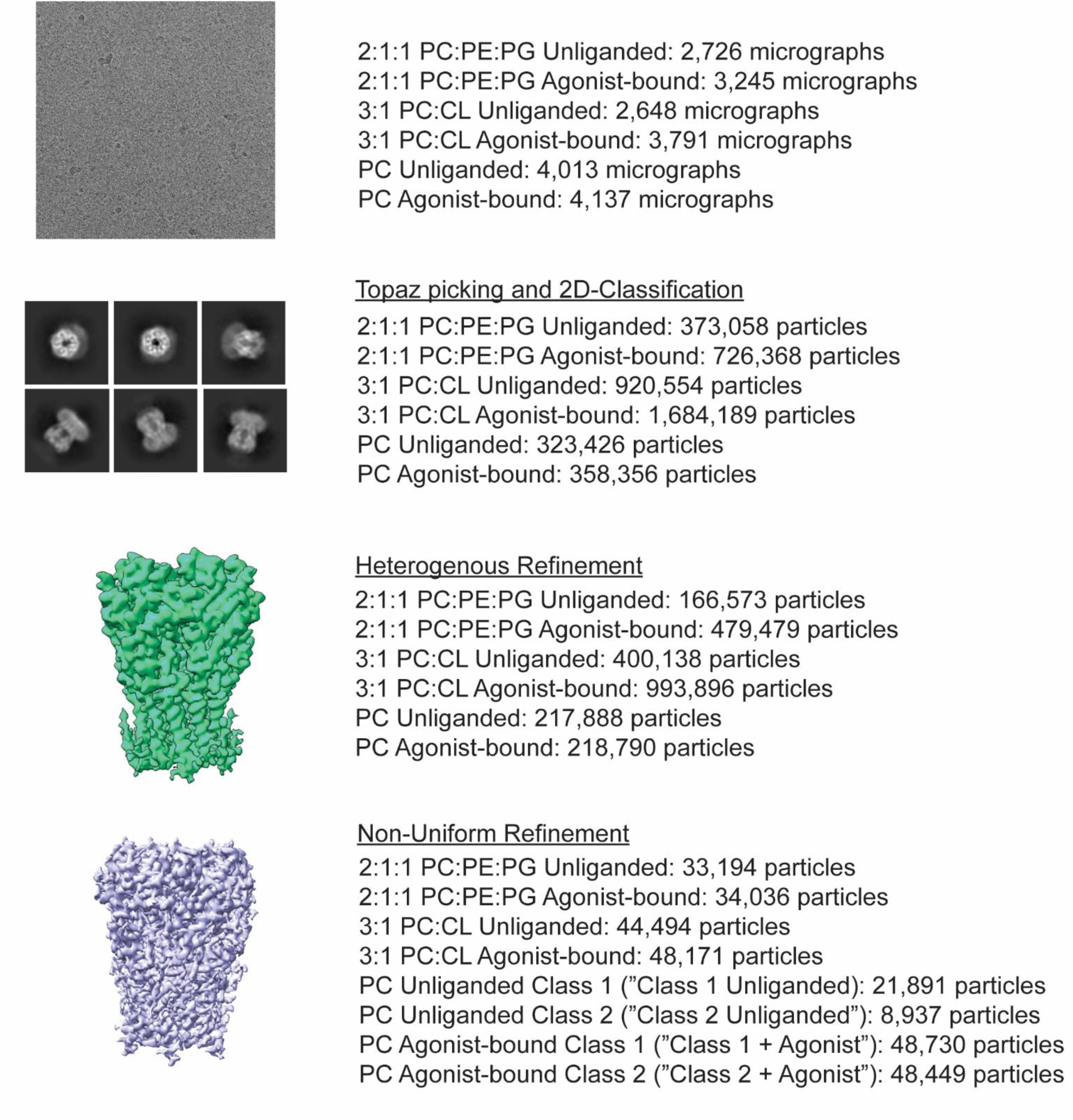
Summary of the single particle cryo-EM analysis in cryoSPARC. Listed are the numbers of micrographs and particles included in each step of the analysis including particle picking, heterogeneous refinement and non-uniform refinement. In the case of the POPC structures, classes 1 and 2 are structures from 3DVA clustering.

**Supplementary Figure 6:**
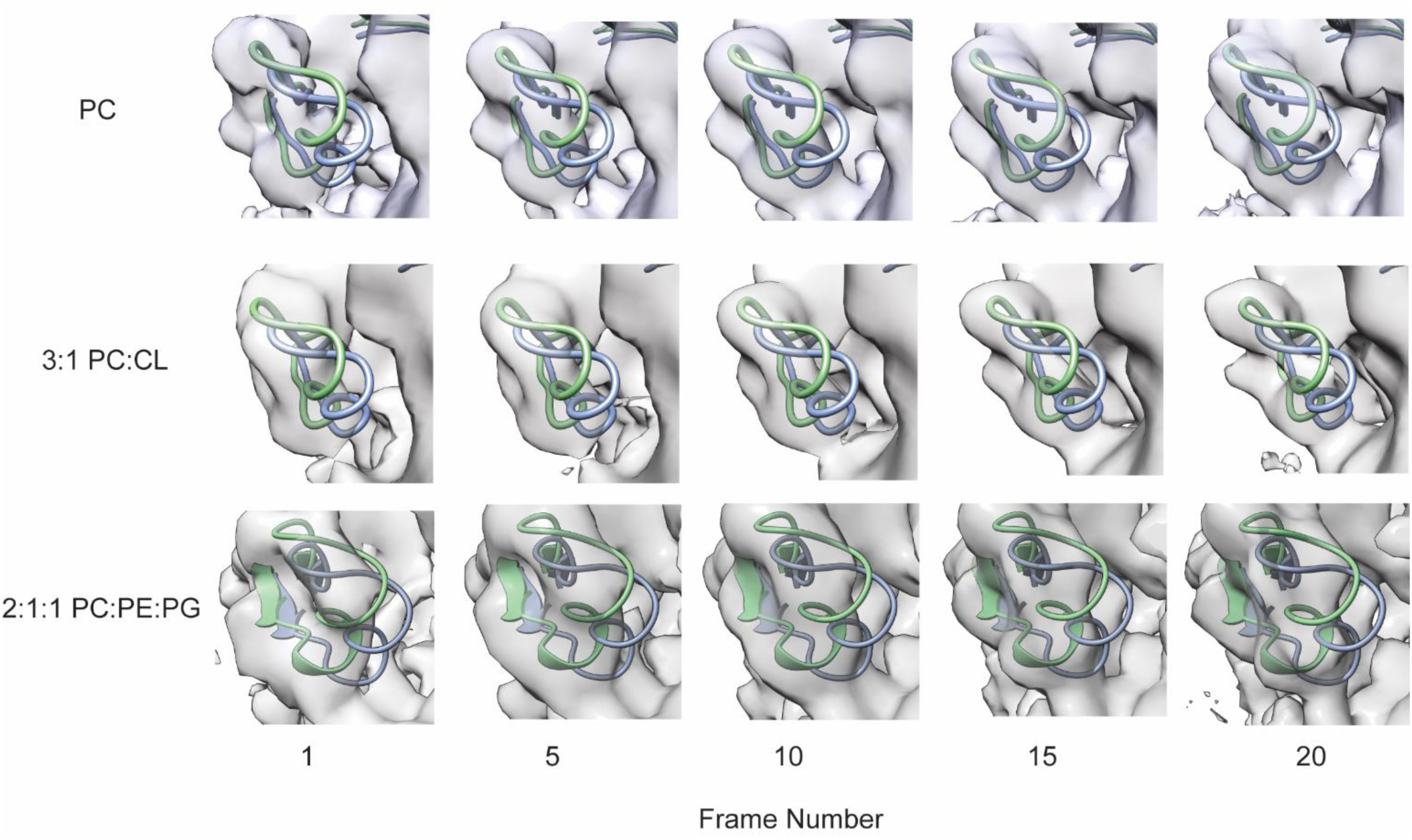
Cryo-EM maps from 3DVA snapshots from ELIC5 unliganded datasets. For each unliganded ELIC5 cryo-EM dataset in a POPC, 3:1 POPC:DPOCL and 2:1:1 POPC:POPE:POPG lipid environment, snapshots of Loop C in the agonist binding site are shown from the 3DVA along the first variability component (i.e. reaction coordinate). For reference, the structural models of ELIC5 unliganded in 2:1:1 POPC:POPE:POPG (green) and ELIC5 agonist-bound in 2:1:1 POPC:POPE:POPG (blue) are shown. The cryo-EM maps show movement in Loop C in the ELIC5 unliganded PC dataset.

**Supplementary Figure 7:**
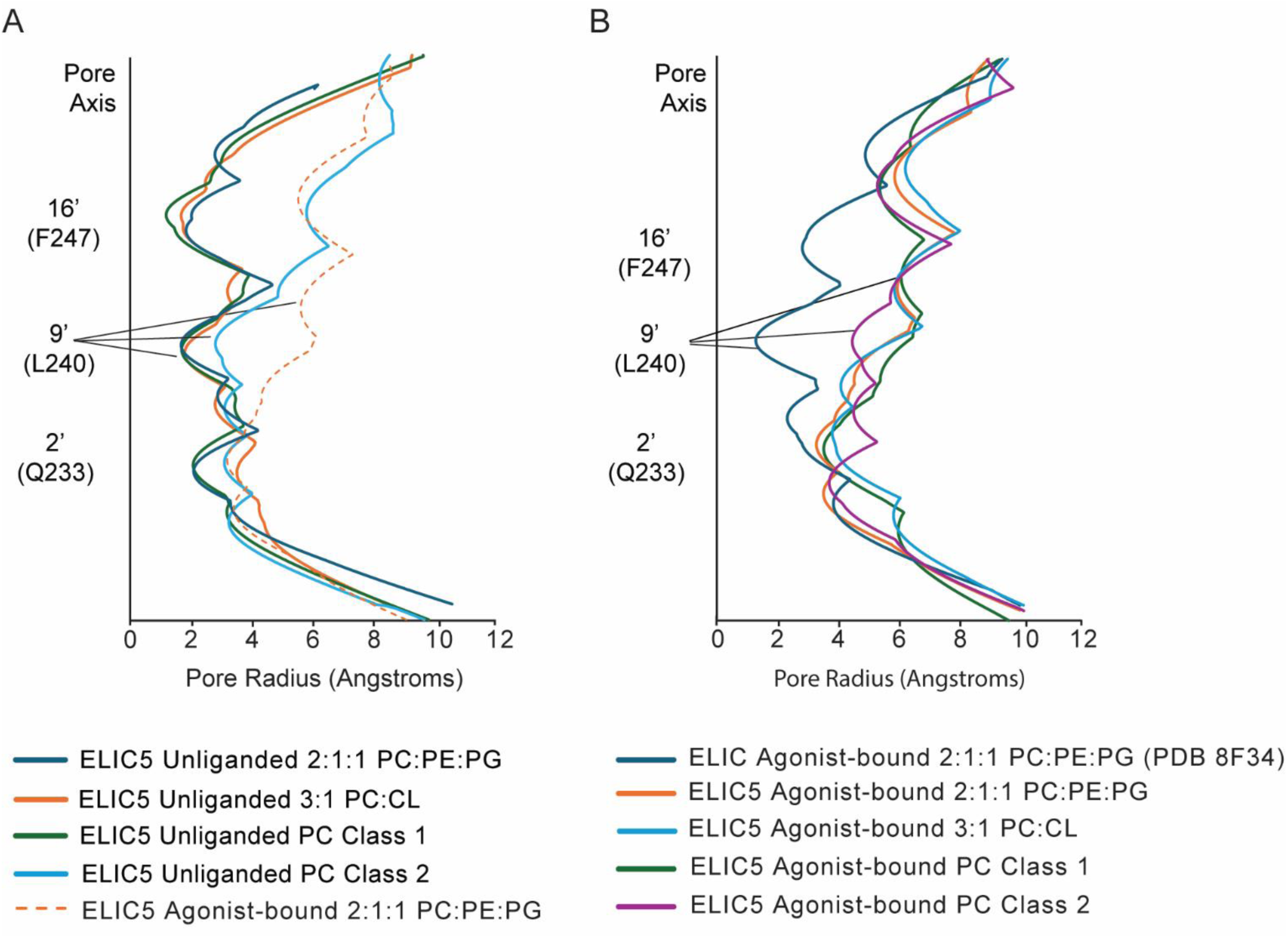
Analysis of pore dimensions in the ELIC5 structures. Plot of pore radius for the indicated structures from this study generated using HOLE. The pore profiles in (A) are unliganded ELIC5 structures with the ELIC5 agonist-bound structure in 2:1:1 POPC:POPE:POPG for comparison. The pore profiles in (B) are the agonist-bound structures with agonist-bound WT ELIC (PDB 8F34) structures for comparison.

**Supplementary Figure 8:**
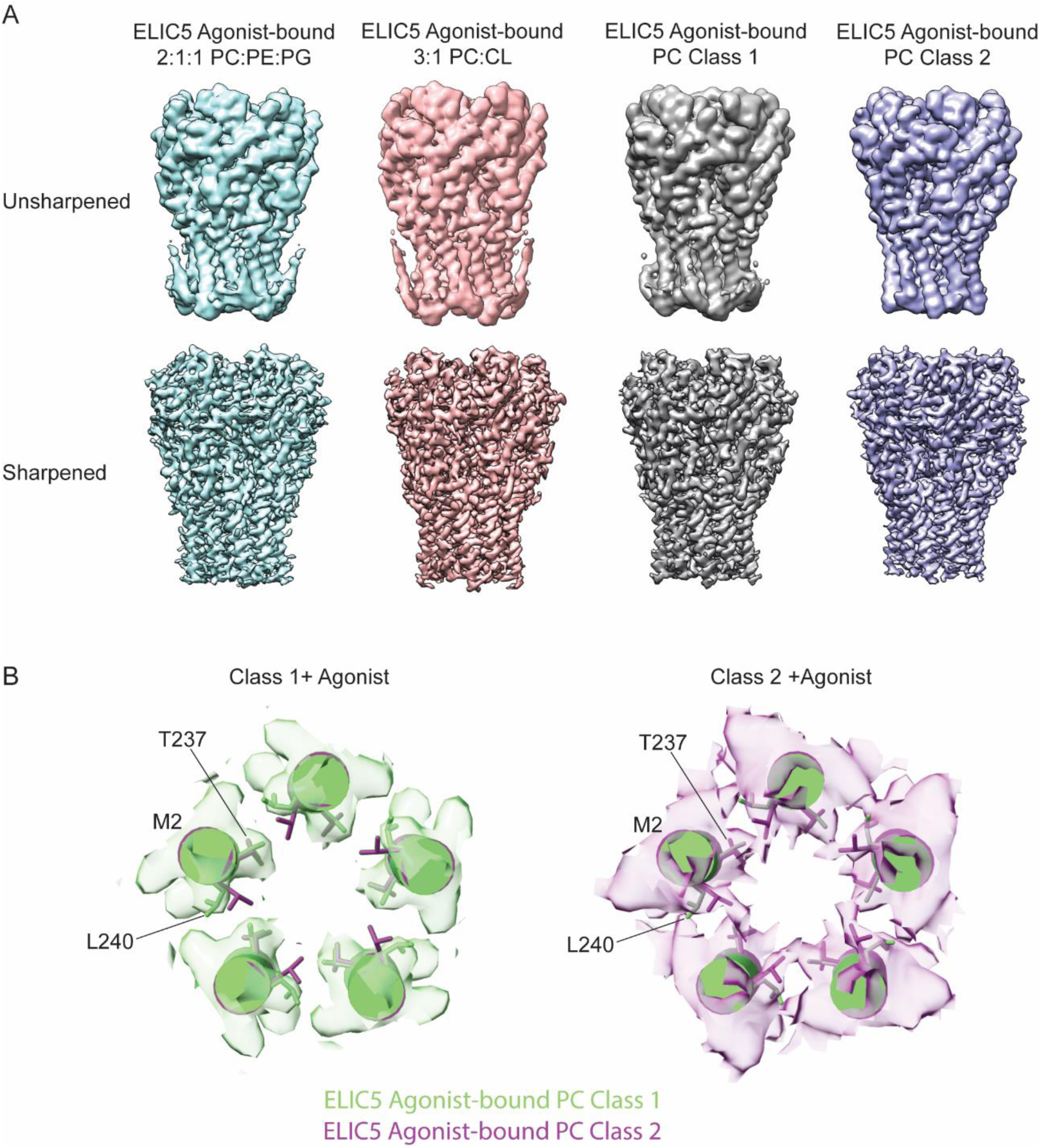
Structural comparison of ELIC5 agonist-bound structures in different lipid environments. (A) Unsharpened and sharpened cryo-EM maps of ELIC5 with agonist in 2:1:1 POPC:POPE:POPG, 3:1 POPC:DPOCL, and POPC lipid environments. The cryo-EM density for M4 is absent in the agonist-bound POPC structures, and therefore this helix was not included in the models. (B) Comparison of “class 1 + agonist” (green) and “class 2 + agonist” (purple) cryo-EM maps for ELIC5 in POPC showing M2 with a view down the pore axis. The models of M2 for the corresponding structures are shown with stick representation of L240 and T237. The structures differ in the side chain orientation of L240.

**Supplementary Figure 9:**
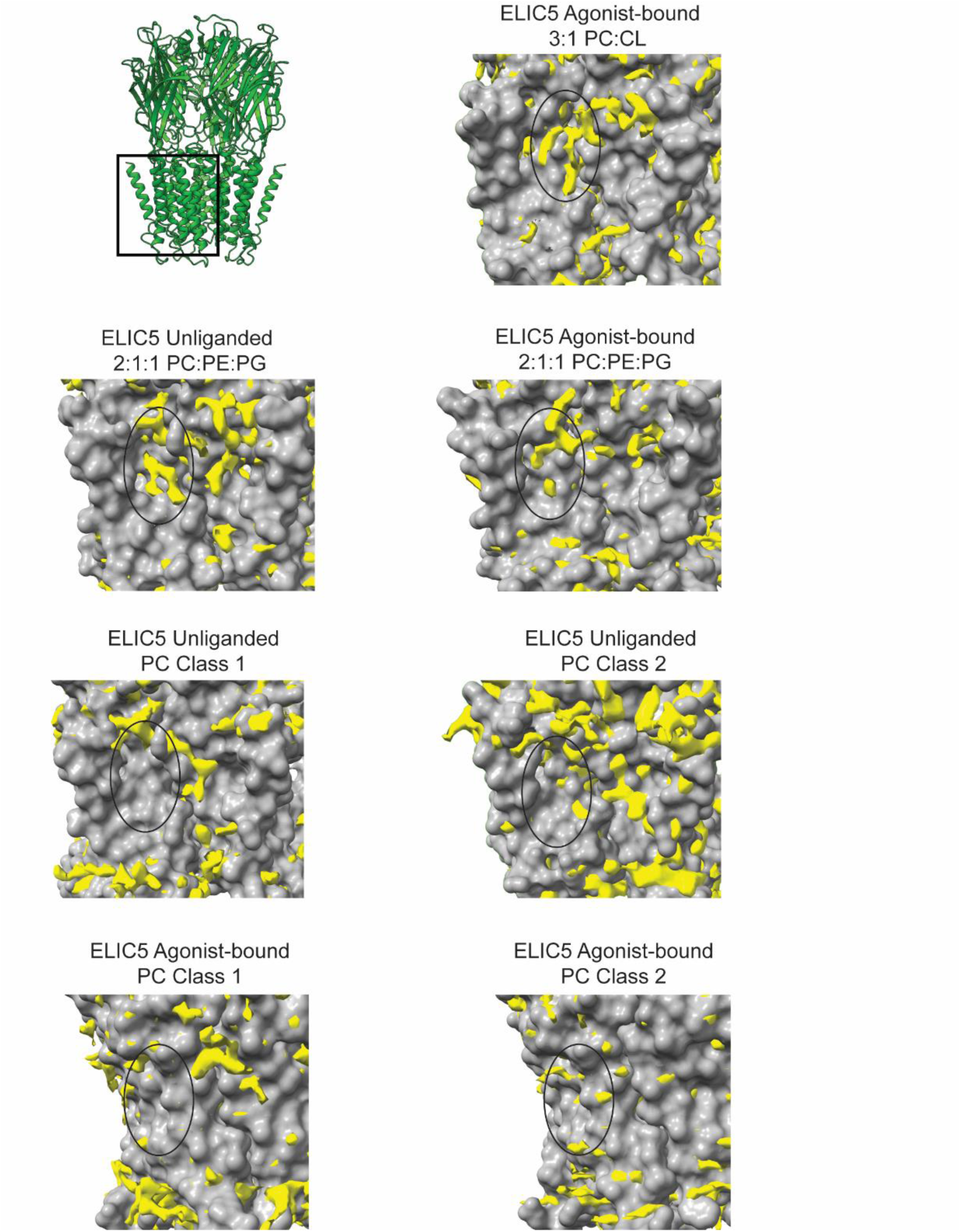
Non-protein densities around the TMD of ELIC structures. For each indicated structure, a surface representation of the protein model is shown of the TMD centered on the M3-M4 groove (indicated by a black circle). The non-protein cryo-EM densities are shown in yellow using the same density threshold for all structures. Weak lipid-like densities are appreciated near M3 and M4 in the 2:1:1 POPC:POPE:POPG and 3:1 POPC:DPOCL structures with the clearest density observed in the unliganded 3:1 POPC:DPOCL structure. Note that M4 was not resolved in the agonist-bound POPC structures so this helix was not included in the models.

**Supplementary Figure 10:**
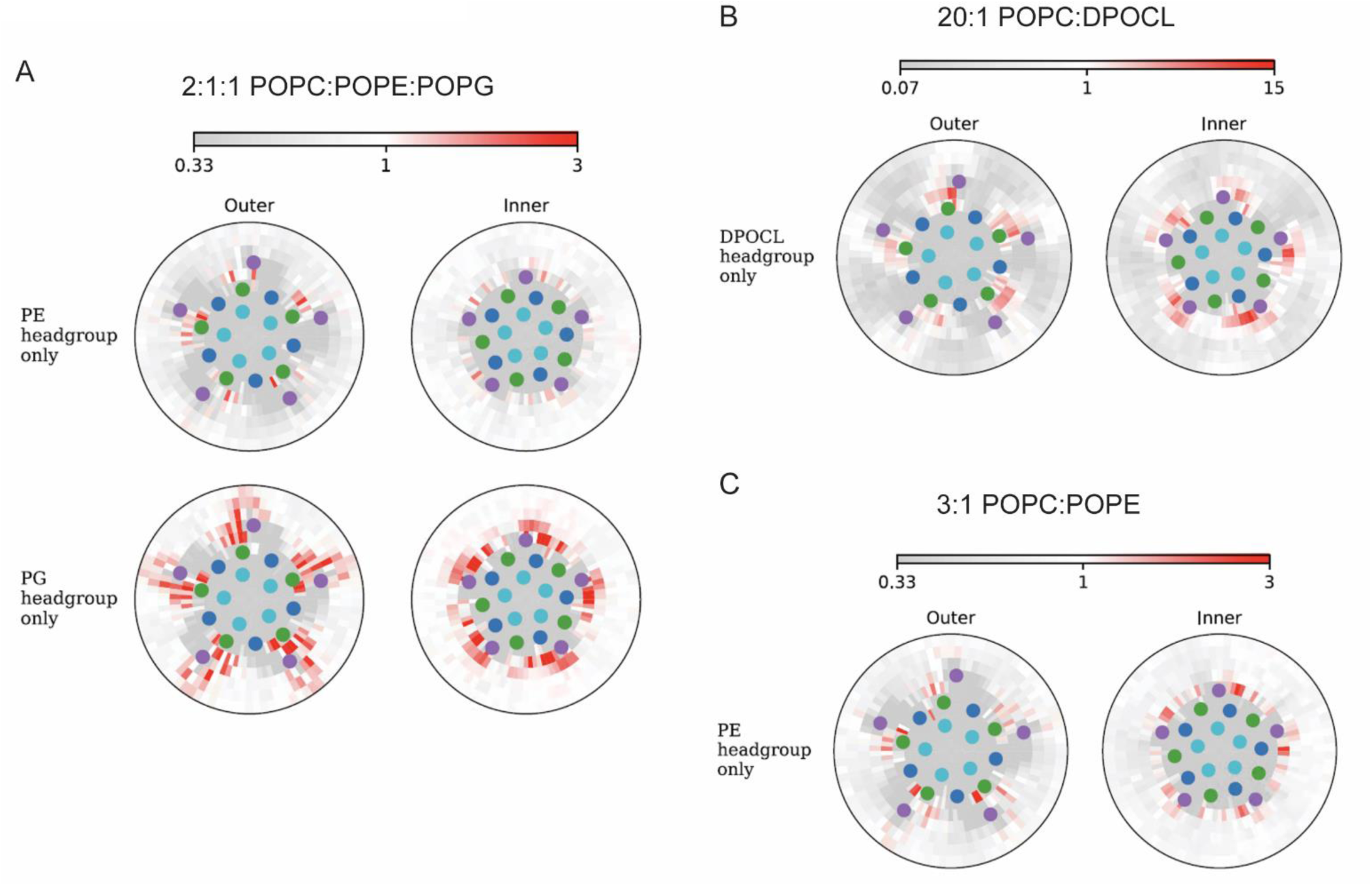
Phospholipid interactions with the agonist-bound ELIC 5 open-channel structure (PDB 9NGC). Radial distribution plots from the CGMD simulation of the ELIC5 open-channel structure in the three indicated lipid environments: (A) 2:1:1 POPC:POPE:POPG in which POPE and POPG density enrichment is shown, (B) 20:1 POPC:DPOCL in which DPOCL density enrichment is shown, and (C) 3:1 POPC:POPE in which POPE density enrichment is shown within 5 nm of the protein center. The analysis was performed with the phospholipid headgroups only. The colored dots represent the center of mass of each TMD helix. Shown are the outer and inner leaflet.

**Supplementary Figure 11:**
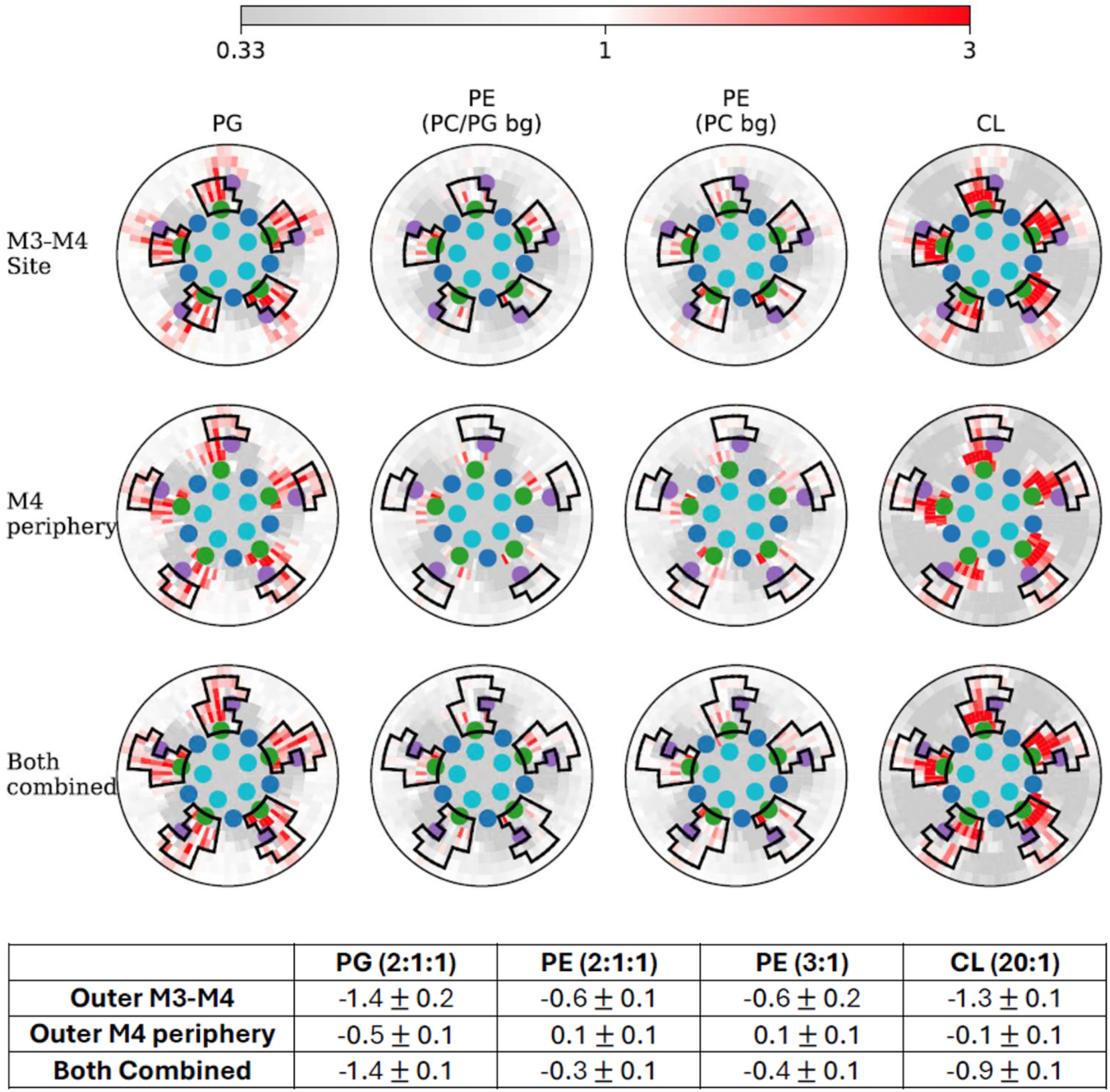
Affinity of phospholipids for the agonist-bound ELIC5 open-channel structure (PDB 9NGC). Radial distribution plots as shown in Supplementary Figure 10 for the outer leaflet. “PG” is the distribution for POPG from a 2:1:1 POPC:POPE:POPG membrane. “PE (PC/PG bg)” is the distribution for POPE from a 2:1:1 POPC:POPE:POPG membrane. “PE (PC bg)” is the distribution for POPE from a 3:1 POPC:POPE membrane. “CL” is the distribution for DPOCL from a 3:1 POPC:DPOCL membrane. The black outlines demarcate the site definitions for the M3-M4 outer leaflet site, an M4 peripheral site, or both combined. The table shows Δ𝐺_bind_ values for the indicated lipids at these sites (± SD).

**Supplementary Figure 12:**
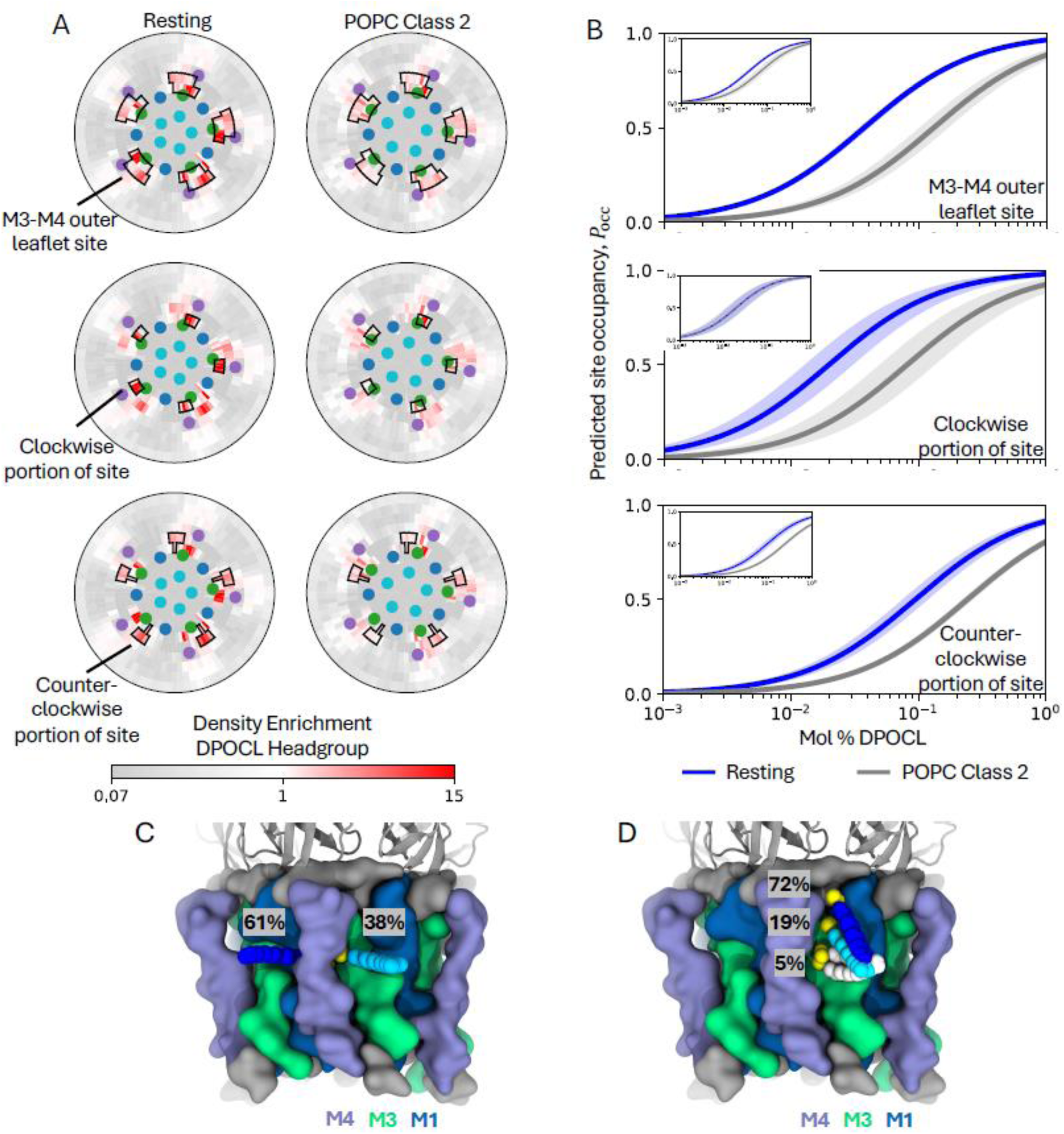
State-dependent binding of DPOCL to an outer leaflet M3-M4 site. (A) Radial distribution plots from the CGMD simulation showing density enrichment of DPOCL headgroups in the outer leaflet within 5 nm of the protein center. The colored dots represent the center of mass of each TMD helix. Black outlines demarcate the site definitions for the M3-M4 outer leaflet site (top), the clockwise (CW) portion of the site (middle), and the counterclockwise (CCW) portion of the site (bottom). (B) Predicted occupancy (𝑃_occ_) of the M3-M4 site (top), CW portion (middle) and CCW portion (bottom), as a function of mol% DPOCL. The shaded region represents 95% confidence interval with n=20 for the resting state and n=15 for POPC class 2, which is the “class 2 unliganded” structure. Insets show 𝑃_occ_ when POPC Class 2 site Δ𝐺_bind_ values are corrected for accessible area. (C-D) Molecular image of the resting state ELIC5 structure from the CGMD simulation with the average positions of most occupied clusters of DPOCL when its headgroup is in the CW portion (C) and CCW portion (D) of the site near M3-M4. The average DPOCL headgroup positions for each cluster are shown as yellow VdW spheres; average tail positions are shown as blue or white VdWspheres. The percentages correspond to the relative frequency of each binding mode.

**Supplementary Figure 13:**
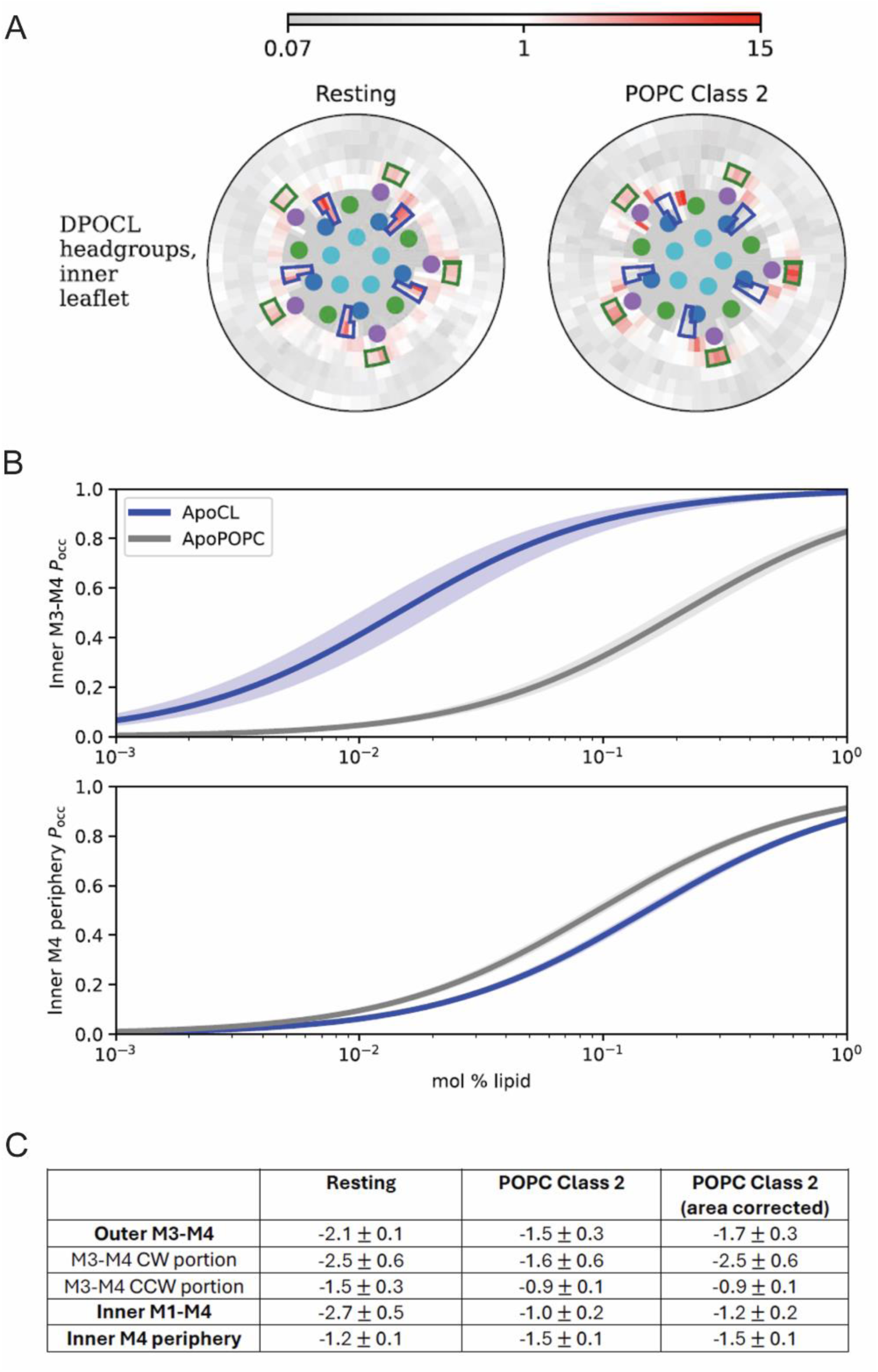
State-dependent binding of DPOCL to an inner leaflet M1-M4 site. (A) Radial distribution plots from the CGMD simulation showing density enrichment of DPOCL headgroups in the inner leaflet within 5 nm of the protein center. The colored dots represent the center of mass of each TMD helix. The outlines demarcate the site definitions for the M1-M4 inner leaflet site and an M4 periphery site in the inner leaflet. (B) Predicted occupancy (𝑃_occ_) of the M1-M4 site (top) and the M4 periphery site (bottom) as a function of mol% DPOCL. The shaded region represents 95% confidence interval with n=20 for the resting state and n=15 for POPC class 2, which is the “class 2 unliganded” structure. (C) Table shows Δ𝐺_bind_ values for DPOCL at all the sites from this figure and Supplementary Figure 12, comparing the resting state structure and POPC class 2, which is the “unliganded class 2” structure (± SD). Also indicated are the Δ𝐺_bind_ values for POPC class 2 when 𝑃_occ_ is corrected for accessible surface area.

**Supplementary Table 1:**
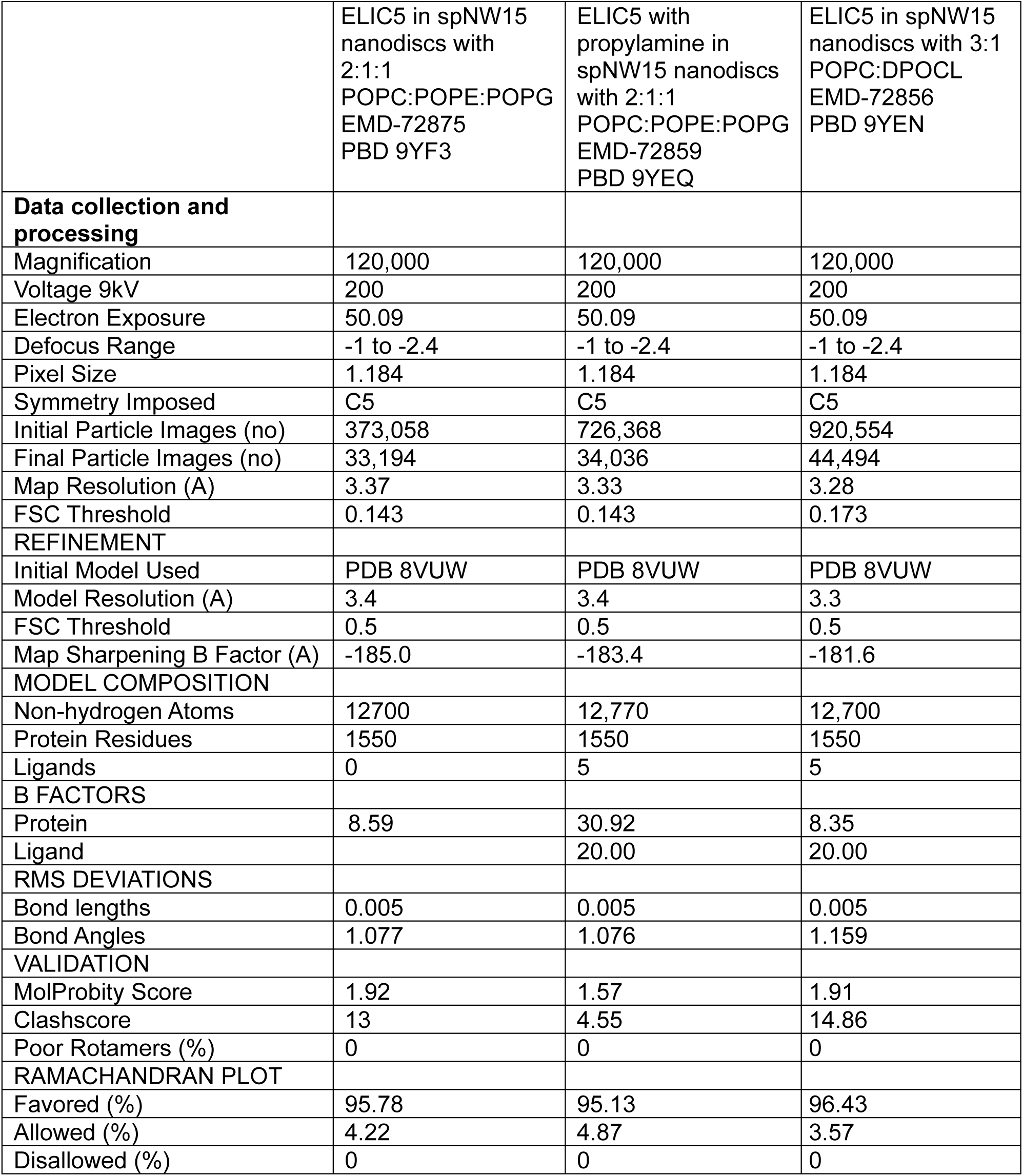
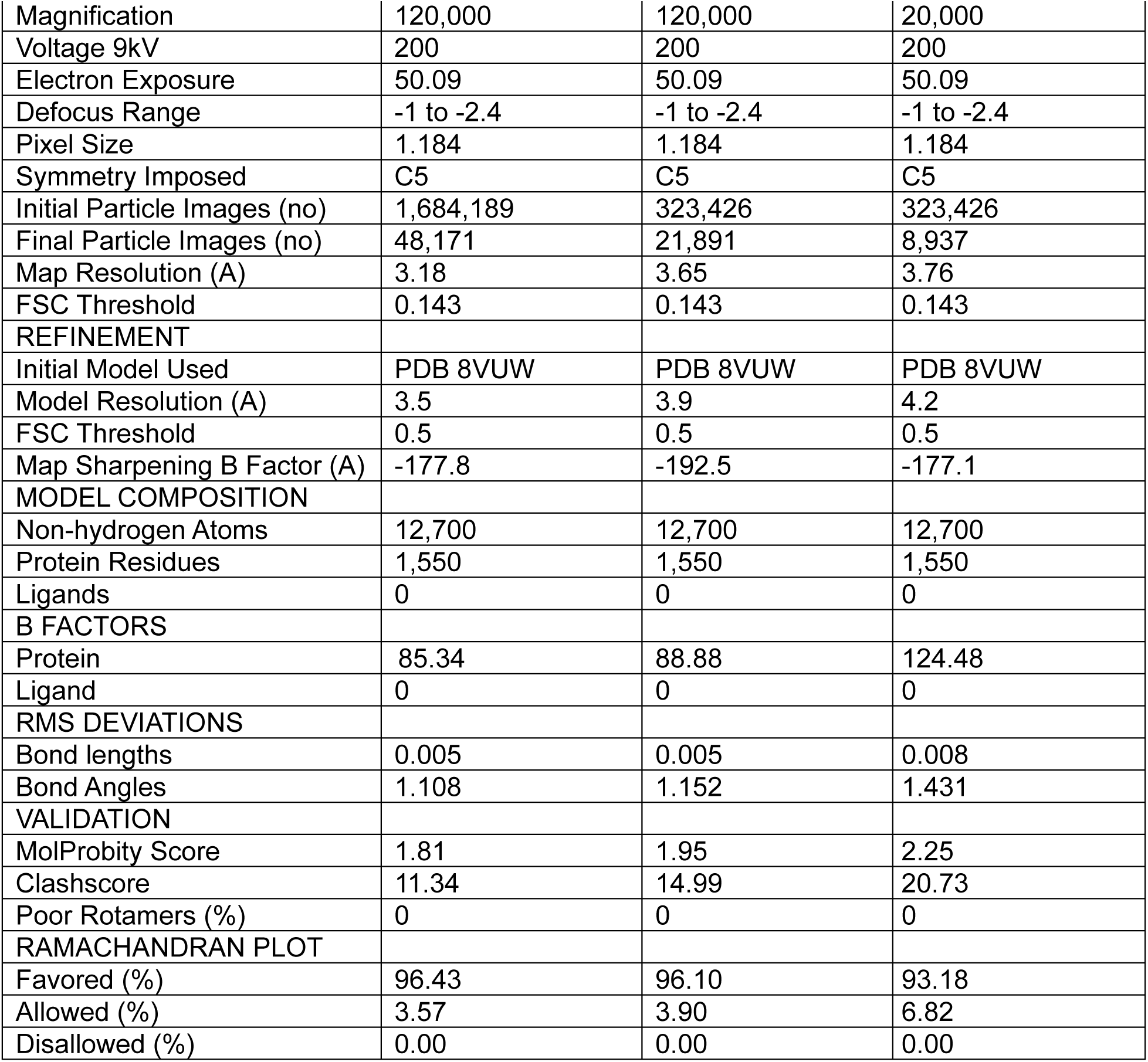
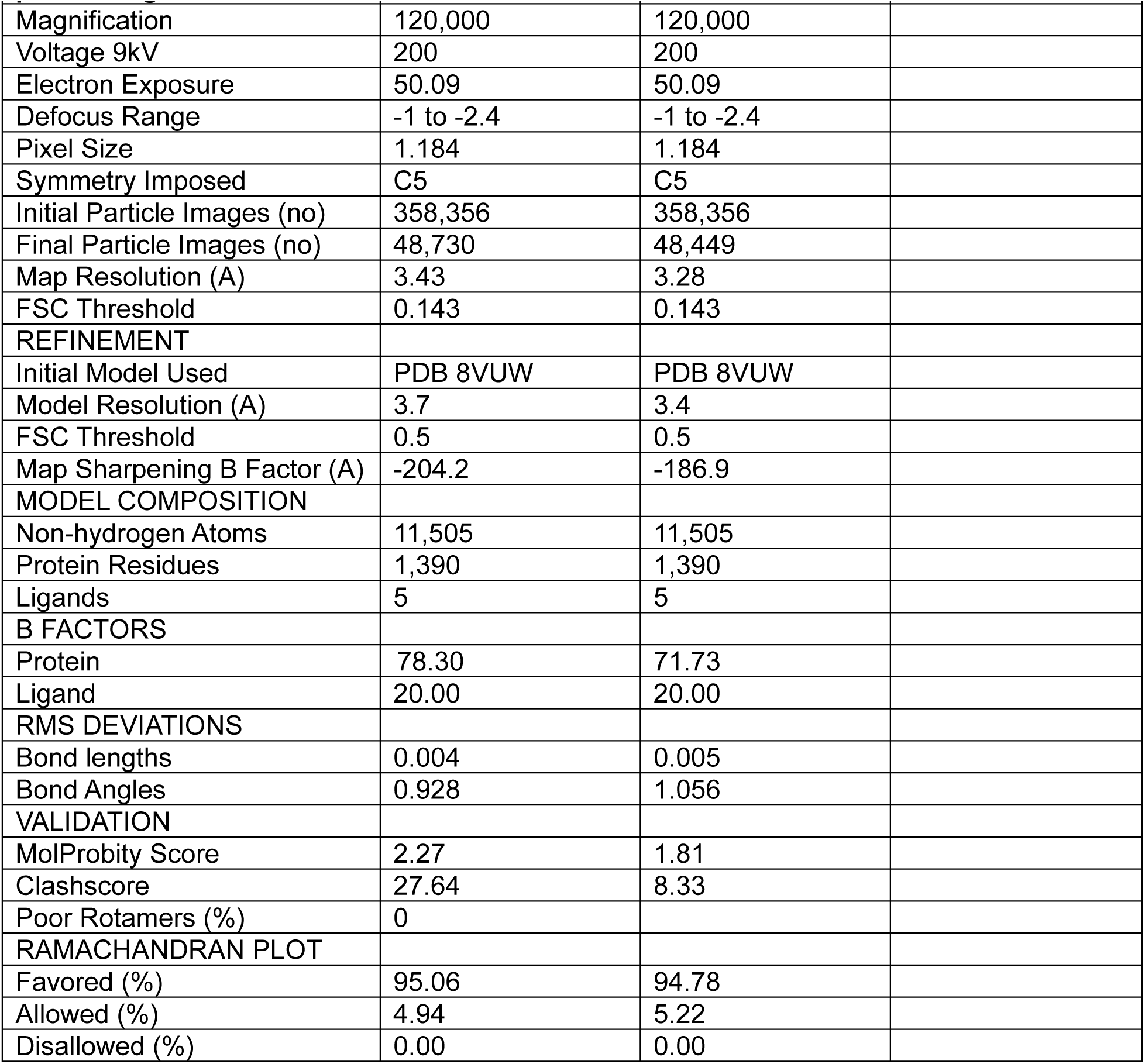
Summary of cryo-EM data collection and refinement parameters

|  |  |  |  |
| --- | --- | --- | --- |
|  | ELIC5 in spNW15 nanodiscs with 2:1:1 POPC:POPE:POPG EMD-72875 PBD 9YF3 | ELIC5 with propylamine in spNW15 nanodiscs with 2:1:1 POPC:POPE:POPG EMD-72859 PBD 9YEQ | ELIC5 in spNW15 nanodiscs with 3:1 POPC:DPOCL EMD-72856 PBD 9YEN |
| **Data collection and processing** |  |  |  |
| Magnification | 120,000 | 120,000 | 120,000 |
| Voltage 9kV | 200 | 200 | 200 |
| Electron Exposure | 50.09 | 50.09 | 50.09 |
| Defocus Range | -1 to -2.4 | -1 to -2.4 | -1 to -2.4 |
| Pixel Size | 1.184 | 1.184 | 1.184 |
| Symmetry Imposed | C5 | C5 | C5 |
| Initial Particle Images (no) | 373,058 | 726,368 | 920,554 |
| Final Particle Images (no) | 33,194 | 34,036 | 44,494 |
| Map Resolution (Å) | 3.37 | 3.33 | 3.28 |
| FSC Threshold | 0.143 | 0.143 | 0.173 |
| **REFINEMENT** |  |  |  |
| Initial Model Used | PDB 8VUW | PDB 8VUW | PDB 8VUW |
| Model Resolution (Å) | 3.4 | 3.4 | 3.3 |
| FSC Threshold | 0.5 | 0.5 | 0.5 |
| Map Sharpening B Factor (Å) | -185.0 | -183.4 | -181.6 |
| **MODEL COMPOSITION** |  |  |  |
| Non-hydrogen Atoms | 12700 | 12,770 | 12,700 |
| Protein Residues | 1550 | 1550 | 1550 |
| Ligands | 0 | 5 | 5 |
| **B FACTORS** |  |  |  |
| Protein | 8.59 | 30.92 | 8.35 |
| Ligand |  | 20.00 | 20.00 |
| **RMS DEVIATIONS** |  |  |  |
| Bond lengths | 0.005 | 0.005 | 0.005 |
| Bond Angles | 1.077 | 1.076 | 1.159 |
| **VALIDATION** |  |  |  |
| MolProbity Score | 1.92 | 1.57 | 1.91 |
| Clashscore | 13 | 4.55 | 14.86 |
| Poor Rotamers (%) | 0 | 0 | 0 |
| **RAMACHANDRAN PLOT** |  |  |  |
| Favored (%) | 95.78 | 95.13 | 96.43 |
| Allowed (%) | 4.22 | 4.87 | 3.57 |
| Disallowed (%) | 0 | 0 | 0 |
|  | ELIC5 with propylamine in spNW15 nanodiscs with 3:1 POPC:DPOCL EMD-72858 PBD 9YEP | ELIC5 in spNW15 nanodiscs with POPC Class 1 EMD-72861 PBD 9YES | ELIC5 in spNW15 nanodiscs with POPC Class 2 EMD-72865 PBD 9YEW |
| **Data collection and processing** |  |  |  |
| Magnification | 120,000 | 120,000 | 20,000 |
| Voltage 9kV | 200 | 200 | 200 |
| Electron Exposure | 50.09 | 50.09 | 50.09 |
| Defocus Range | -1 to -2.4 | -1 to -2.4 | -1 to -2.4 |
| Pixel Size | 1.184 | 1.184 | 1.184 |
| Symmetry Imposed | C5 | C5 | C5 |
| Initial Particle Images (no) | 1,684,189 | 323,426 | 323,426 |
| Final Particle Images (no) | 48,171 | 21,891 | 8,937 |
| Map Resolution (Å) | 3.18 | 3.65 | 3.76 |
| FSC Threshold | 0.143 | 0.143 | 0.143 |
| REFINEMENT |  |  |  |
| Initial Model Used | PDB 8VUW | PDB 8VUW | PDB 8VUW |
| Model Resolution (Å) | 3.5 | 3.9 | 4.2 |
| FSC Threshold | 0.5 | 0.5 | 0.5 |
| Map Sharpening B Factor (Å) | -177.8 | -192.5 | -177.1 |
| MODEL COMPOSITION |  |  |  |
| Non-hydrogen Atoms | 12,700 | 12,700 | 12,700 |
| Protein Residues | 1,550 | 1,550 | 1,550 |
| Ligands | 0 | 0 | 0 |
| B FACTORS |  |  |  |
| Protein | 85.34 | 88.88 | 124.48 |
| Ligand | 0 | 0 | 0 |
| RMS DEVIATIONS |  |  |  |
| Bond lengths | 0.005 | 0.005 | 0.008 |
| Bond Angles | 1.108 | 1.152 | 1.431 |
| VALIDATION |  |  |  |
| MolProbity Score | 1.81 | 1.95 | 2.25 |
| Clashscore | 11.34 | 14.99 | 20.73 |
| Poor Rotamers (%) | 0 | 0 | 0 |
| RAMACHANDRAN PLOT |  |  |  |
| Favored (%) | 96.43 | 96.10 | 93.18 |
| Allowed (%) | 3.57 | 3.90 | 6.82 |
| Disallowed (%) | 0.00 | 0.00 | 0.00 |

|  |  |  |
| --- | --- | --- |
|  | ELIC5 with<br>propylamine in<br>spNW15 nanodiscs<br>with POPC Class 1<br>EMD-72867<br>PBD 9YEY | ELIC5 with<br>propylamine in<br>spNW15<br>nanodiscs with<br>POPC Class 2<br>EMD-72869<br>PBD 9YF1 |
| **Data collection and**<br>**processing** |  |  |
| Magnification | 120,000 | 120,000 |
| Voltage 9kV | 200 | 200 |
| Electron Exposure | 50.09 | 50.09 |
| Defocus Range | -1 to -2.4 | -1 to -2.4 |
| Pixel Size | 1.184 | 1.184 |
| Symmetry Imposed | C5 | C5 |
| Initial Particle Images (no) | 358,356 | 358,356 |
| Final Particle Images (no) | 48,730 | 48,449 |
| Map Resolution (Å) | 3.43 | 3.28 |
| FSC Threshold | 0.143 | 0.143 |
| REFINEMENT |  |  |
| Initial Model Used | PDB 8VUW | PDB 8VUW |
| Model Resolution (Å) | 3.7 | 3.4 |
| FSC Threshold | 0.5 | 0.5 |
| Map Sharpening B Factor (Å) | -204.2 | -186.9 |
| MODEL COMPOSITION |  |  |
| Non-hydrogen Atoms | 11,505 | 11,505 |
| Protein Residues | 1,390 | 1,390 |
| Ligands | 5 | 5 |
| B FACTORS |  |  |
| Protein | 78.30 | 71.73 |
| Ligand | 20.00 | 20.00 |
| RMS DEVIATIONS |  |  |
| Bond lengths | 0.004 | 0.005 |
| Bond Angles | 0.928 | 1.056 |
| VALIDATION |  |  |
| MolProbity Score | 2.27 | 1.81 |
| Clashscore | 27.64 | 8.33 |
| Poor Rotamers (%) | 0 |  |
| RAMACHANDRAN PLOT |  |  |
| Favored (%) | 95.06 | 94.78 |
| Allowed (%) | 4.94 | 5.22 |
| Disallowed (%) | 0.00 | 0.00 |

**<u>Supplementary Video 1:</u>** The 3DVA reaction coordinate along the first variability component for the unliganded ELIC5 structure in POPC nanodiscs. The video shows a side view of the full ion channel.

**<u>Supplementary Video 2:</u>** The 3DVA reaction coordinate along the first variability component for the unliganded ELIC5 structure in POPC nanodiscs. The video shows a top-down view along the pore axis of the ion channel from the extracellular side.

## Notes

### Competing Interest Statement

The authors have declared no competing interest.

